# tiRNA signaling via stress-regulated vesicle transfer in the hematopoietic niche

**DOI:** 10.1101/2021.04.13.439696

**Authors:** Youmna S. Kfoury, Fei Ji, Michael Mazzola, David B. Sykes, Allison K. Scherer, Anthony Anselmo, Yasutoshi Akiyama, Francois Mercier, Nicolas Severe, Konstantinos D. Kokkaliaris, Thomas Brouse, Borja Saez, Jefferson Seidl, Ani Papazian, Pavel Ivanov, Michael K. Mansour, Ruslan I. Sadreyev, David T. Scadden

## Abstract

Extracellular vesicles transfer complex biologic material between cells, whose role in *in-vivo* organismal physiology is poorly defined. Here, we demonstrate that osteoblastic cells in the bone marrow elaborate extracellular vesicles that are taken up by hematopoietic progenitor cells *in vivo*. Genotoxic or infectious stress rapidly increased stromal-derived extracellular vesicle transfer to granulocyte-monocyte progenitors. Stimulating osteoblastic cells with parathyroid hormone or activating its receptor enhanced extracellular vesicle transfer, myeloid recovery post radiation and improved animal survival from *Candida* sepsis. The extracellular vesicles contained tiRNAs known to modulate protein translation. 5’-ti-Pro-CGG-1 was preferentially abundant in osteoblast-derived extracellular vesicles and when transferred to granulocyte macrophage progenitors, increased protein translation, cell proliferation and myeloid differentiation. Therefore, EV-mediated tiRNA transfer provides a stress modulated signaling axis distinct from conventional cytokine-driven stress responses.

**One sentence summary:** Stress regulated tiRNA transfer alters hematopoiesis

## Introduction

Stem cell niches, are specialized local microenvironments that modulate stem and progenitor populations of a tissue (Jones and Wagers, 2008). They have largely been defined in terms of the cells comprising them and the cytokines or adhesion molecules produced by them. We hypothesized that interactions between mesenchymal stromal cells of a niche and the stem/progenitor populations with which they interact may have a broader range of modulators. Specifically, we considered it highly likely that niche-stem cell interactions evolved early in metazoans, perhaps earlier than the more evolved ligand-receptor interactions that currently dominate our mechanistic understanding of niches.

Extracellular vesicle production is an evolutionary conserved process executed by almost every cell type at steady state or upon activation (van Niel et al., 2018). Originally described as a trash bin for cellular waste (Johnstone et al., 1987), extracellular vesicles (EVs) are conceptually extending to serving as a diagnostic and prognostic tool (Chen et al., 2018; Melo et al., 2015), a method for the delivery of therapeutic molecules (Conlan et al., 2017; Stranford and Leonard, 2017) and as a mechanism for cellular communication (Mathieu et al., 2019). Extracellular vesicles (EVs) are membrane bound and transfer bioactive molecules that alter the physiology of recipient cells. Among these are proteins, lipids and nucleic acids, including mRNA, DNA and small non-coding RNA (sncRNA) including miRNA, tRNA, snoRNA and piRNA (Jeppesen et al., 2019; Tkach and Thery, 2016; Valadi et al., 2007; Wei et al., 2017). Often noted as products of primary cells and cancer cell lines *in vitro*, evidence for a physiological role *in vivo* remains limited to few systems including endothelial cells (Crewe et al., 2018) and adipose tissue (Ying et al., 2017).

It had been noted that mesenchymal cells are producers of extracellular vesicles (EV) and that mesenchymal cells in culture could affect co-cultured hematopoietic stem/progenitors (HSPC) in vitro (Goloviznina et al., 2016; Morhayim et al., 2016; Wen et al., 2016b). We evaluated whether such a process occurs *in vivo*. Defining that it does, we assessed the cargo that was transferred. Notably, we found multiple sncRNA species in EVs, some of which were increased in EV-receiving HSPC. Among these, processed tRNAs (tiRNA) were shown to alter HSPC proliferation. That sncRNA could serve to modulate hematopoiesis was particularly intriguing because we had previously noted that deletion of a RNA processing enzyme, Dicer1, specifically in early osteolineage cells resulted in a dysplastic hematopoietic phenotype . This sometimes resulted in emergence of acute myeloid leukemia, but the hematopoietic cells had intact *Dicer1*. It was not clear how a cell non-autonomous effect could be induced by RNA processing (Raaijmakers et al., 2010). Similarly, we had noted that production of a tRNA processing enzyme, angiopoietin1, Ang1, resulted in processed tRNA in HSPC affecting their quiescence by modifying protein translation (Goncalves et al., 2016; Silberstein et al., 2016). The findings below indicate that specific stromal cells transfer a species of processed tRNA directly to hematopoietic cells through EV creating a cell communication schema distinct from the ligand-receptor paradigm. This signaling process is one that is increased under physiologic stress and may represent a distinctive, perhaps ancient form of niche regulation.

## Results

### Extracellular vesicles shuttle proteins and RNA from osteoblastic to hematopoietic cells in the BM

Exchange of cellular material between BMMS and HSPCs was evaluated using mouse reporter models with GFP or GFP^Topaz^ expressed under control of promoters active in specific mesenchymal cells that are known extrinsic regulators of HSPCs (Fig. 1A) (Morrison and Scadden, 2014). Osteocalcin GFP-Topaz (Bilic-Curcic et al., 2005) (Ocn-GFP^Topaz^) and Collagen 1-GFP (Kalajzic et al., 2003) (Col1-GFP) marked osteoblastic cells, Osterix-Cre::GFP (Rodda and McMahon, 2006) (Osx-GFP) marked osteoprogenitor cells and Nestin-GFP (Mignone et al., 2004) (Nes-GFP) marked primitive mesenchymal stromal cells (MSC). GFP is 27kDa, prohibiting its intercellular transfer through gap junctions (upper limit, 1kDa) (Nielsen et al., 2012).

**Fig. 1:**
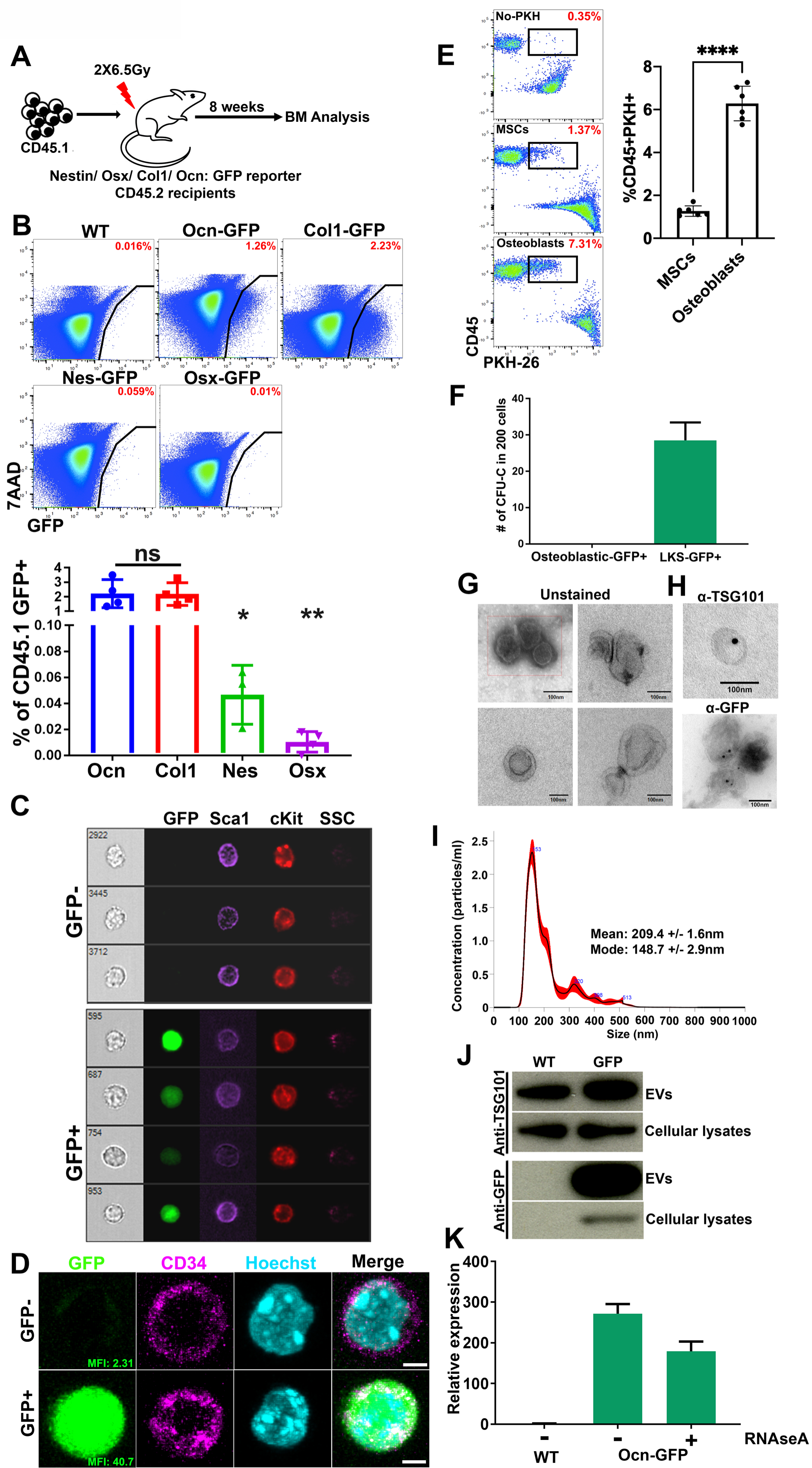
Select mesenchymal subsets are major sources of EVs in the BM. **(A)** To investigate the transfer of stromal derived sncRNAs via EVs we transplanted lethally irradiated reporter mice that express GFP in specific mesenchymal subsets at different stages of differentiation. **(B)** Frequency of GFP+ cells in donor CD45.1+ BM (parent gate). Data represent independent biological replicates. **(C)** Imaging flow cytometry **(D)** confocal imaging on sorted LKS^GFP+/-^ cells from Ocn-GFP^Topaz^ animals. Scale bar = 3µm. **(E)** Differential transfer of PKH-26 labeled EVs from MSCs or osteoblasts to co-cultured GMPs as shown by flow cytometry. Gates are on live cells. **(F)** Methyl cellulose colonies comparing osteoblasts to LKS^GFP+,^ n=2. **(G)** Transmission electron microscopy of BM derived EVs. **(H)** Immunogold staining using 15nm gold beads (TSG-101), 10nm gold beads (GFP), Scale bars in G and H = 100nm. **(I)** NTA showing size distribution of EVs isolated from the mouse BM. The mean and mode are calculated based on 5 measurements. **(J)** Western blot analysis for TSG101 and GFP on EVs and cellular lysates. **(K)** GFP qPCR on RNA extracted from RNAse A treated EVs. Data represent three technical replicates. Data in B and E is presented as mean ± s.d. *p<0.05, **p<0.01, ****p<0.0001.

Mice were transplanted with wild-type (WT) congenic CD45.1 BM following lethal irradiation. After eight weeks, transplanted BM cells were assessed for the presence of GFP (Fig. 1A). CD45.1 GFP+ cells were ∼40-fold more abundant in Ocn-GFP^Topaz^ and Col1-GFP mice than in Nes-GFP or Osx-GFP recipients (Fig. 1B, fig. S1A). The frequency of GFP+ mesenchymal cells did not correlate with GFP-labeling of hematopoietic cells (fig. S1B). To rule out the effect of radiation, we demonstrated in Ocn-GFP^Topaz^ mice that GFP-labeling within the hematopoietic compartment was comparable in transplanted and non-transplanted animals (fig. S1C). Evident cytoplasmic GFP signal in single hematopoietic cells was observed by imaging flow cytometry ruling out the possibility of osteoblasts in doublets contributing to the signal (Fig. 1C, fig. S1D). Confocal microscopy confirmed that GFP was cytoplasmic rather than non-specifically membrane-bound (Fig. 1D, fig. S1D). We confirmed the enhanced production of EVs by osteoblasts in a co-culture system of PKH-26 labeled osteoblasts or mesenchymal stromal cells (MSCs) with primary hematopoietic progenitors. Osteoblasts labeled 3 times more GMPs compared to MSCs (Fig. 1E). Furthermore, the hematopoietic origin of the GFP+ cells was confirmed using a colony-formation assay: GFP+, Lin-ckit+Sca1+ (LKS) formed GFP-colonies in methylcellulose in contrast to the GFP+CD45-osteoblastic cells from the same animals which did not form any colonies under hematopoietic cell culture conditions (Fig. 1F, fig. S1E-H).

To investigate the transfer of GFP via EVs, we performed transmission electron microscopy analysis (TEM) which revealed cup-shaped, membrane-bound vesicles in Ocn-GFP^Topaz^ BM derived EVs (Fig. 1G). Nanoparticle tracking analysis revealed vesicles with a mean size of 209.4nm (+/- 1.6) and a mode of 148.7nm (+/- 2.9), in keeping with exosome dimensions (Fig. 1I) (Mathieu et al., 2019; van Niel et al., 2018). The exosome specific protein TSG101 was present on the EVs as confirmed by TEM (immunogold staining) and Western blotting (WB) (Fig. 1H, J). GFP was similarly detected in EV preparations by TEM and WB at the protein level (Fig. 1H, J). Additionally, GFP mRNA was detected by qPCR in RNAseA treated Ocn-GFP^Topaz^ BM EVs which was transferred to primary *ex-vivo* cultured GMPs (Fig. 1K, fig. S1J). Finally, the exosome-defining tetraspanins, CD81 and CD9, were evident on the surface of BM EVs by flow cytometry (fig. S1I). Together, these findings demonstrate that among BMMS, osteoblasts are producers of EVs of endocytic origin that transfer GFP protein and mRNA to hematopoietic cells *in vivo*.

### Granulocyte macrophage progenitors are the most abundant EV recipients among HSPCs

Given the role of BMMS in the regulation of HSPC function (Kfoury and Scadden, 2015) and the experimental evidence demonstrating that alteration of specific BMMS results in myeloid malignancies (Dong et al., 2016; Kode et al., 2014; Raaijmakers et al., 2010), we hypothesized that BMMS-derived EVs might regulate HSPCs. Using uptake of GFP as an indicator for EV uptake, we examined HSPC populations: LKS, Lin-cKit+Sca1-CD34+CD16/32lo common myeloid progenitors (CMP), Lin-cKit+Sca1-CD34+CD16/32hi granulocyte macrophage progenitors (GMP), Lin-cKit+Sca1-CD34-CD16/32lo megakaryocyte erythroid progenitors (MEP) and Lin-IL7R+cKit+Sca1+ common lymphoid progenitors (CLP) in the BM of the Ocn-GFP^Topaz^ mice by flow cytometry. GMPs and LKS were labeled at a comparable frequency which was significantly higher than CMPs, MEPs and CLPs (Fig. 2A, fig. S2A). However, the higher frequency of GMPs (0.95% ± 0.15) compared to LKS (0.28% ± 0.05) in BM mononuclear cells results in very low numbers of labeled LKS and significantly higher numbers of labeled GMPs. Labeling of Lin-, cKit+, Sca1+, CD150+, CD48-(LT-HSC) was negligible (fig. S2B). Higher level, but similarly distributed EV uptake, was observed using the Col1-GFP mouse model (fig. S2C-D). However, given that the Col1-GFP model labels a wider population of osteoblasts and pre-osteoblasts in addition to the specificity of the Ocn-GFP^Topaz^ to matrix forming osteoblasts (Bilic-Curcic et al., 2005), we chose to proceed with the latter model for follow up experiments to ensure we are analyzing a homogeneous population of EVs. Imaging flow cytometry and confocal microscopy confirmed single cell GMP^GFP+^ cells and cytoplasmic GFP (Fig. 2B-C). Scatter properties and Wright-Giemsa staining did not reveal any morphological differences between GFP+ GMPs (GMP^GFP+^) and GFP-GMPs (GMP^GFP-^) (Fig. 2D-E). However, GMP^GFP+^ were enriched in colony forming unit capacity with comparable colony size (Fig. 2F, fig. S2F). Among Lin+ cells in the BM, CD11b+ myeloid cells had the highest frequency of labeling (fig. S2G-H). The transfer of EVs between osteoblasts and GMPs was confirmed by confocal imaging of a co-culture between PKH-26 labeled osteoblasts and GMPs isolated from CAG-ECFP animals (Fig. 2G, fig.S2E). White arrows point towards PKH-26 labeled vesicles in GMPs transferred from osteoblasts (Fig. 2G). These vesicles were not detected in GMPs in the absence of labeled osteoblasts (fig. S2E). Using a similar co-culture system, we confirmed the preferential uptake of osteoblast derived EVs by GMPs compared to CMPs (Fig. 2H).

**Fig. 2:**
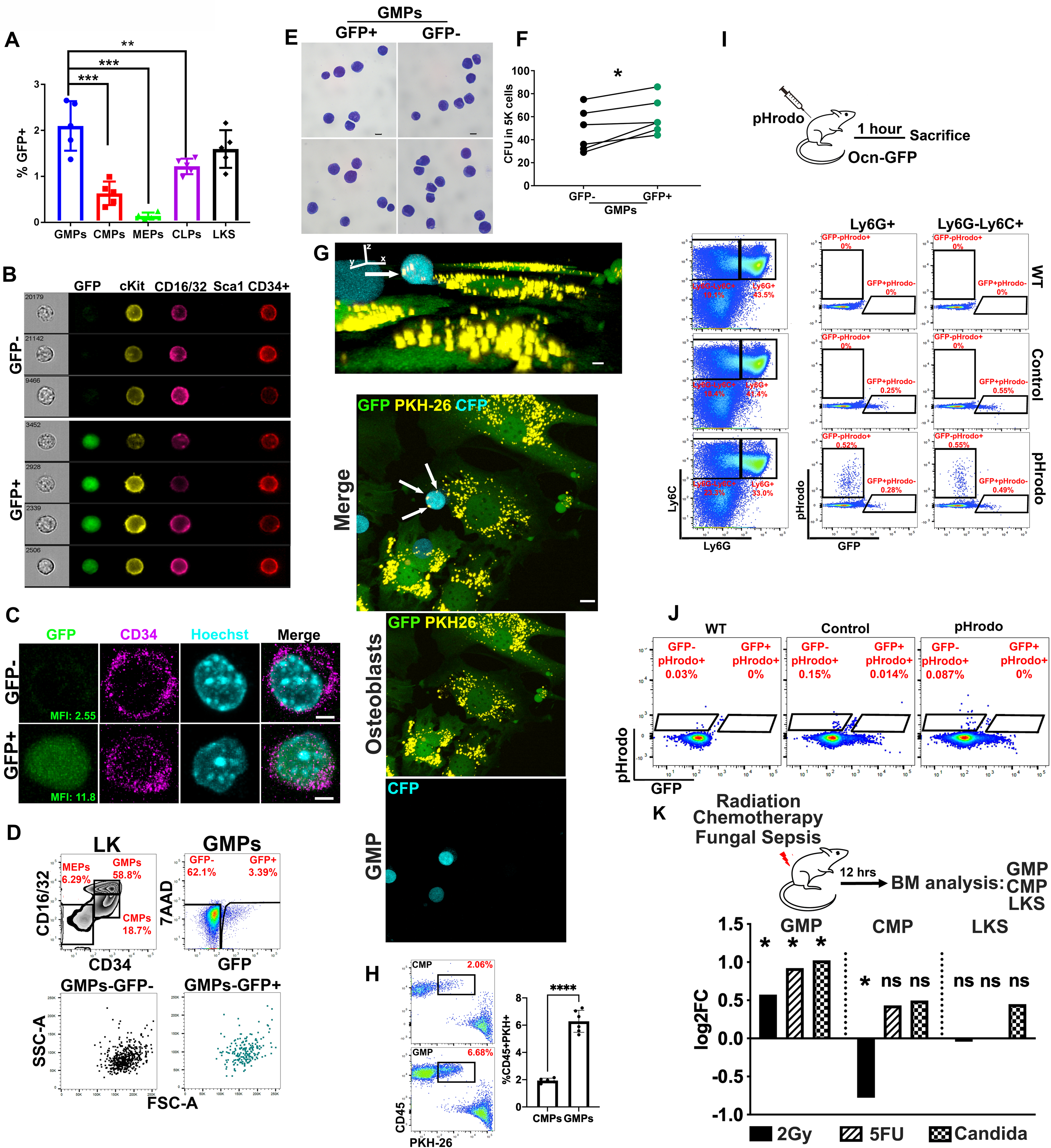
GMPs are major recipients of osteoblastic EVs within the Lin-compartment of the BM. **(A)** Frequency of GFP+ cells (of parent gate) within BM HSPCs. Data represent independent biological replicates. **(B)** Imaging flow cytometry **(C)** confocal imaging of sorted GMP^GFP+^ and GMP^GFP-^. Scale bar = 3µm. **D, E** morphological assessment of GMP^GFP+^ by **(D)** flow cytometry **(E)** and bright field microscopy of Wright Giemsa staining. Scale bar = 10µm. **(F)** Hematopoietic colonies in methyl cellulose comparing GMP^GFP+^ to GMP^GFP-^. Statistical significance is calculated using paired student t-test, * *P*<0.05. Data represents one out of three independent experiments. **(G)** Upper: XYZ view of GMP (CFP) cell with PKH-26 labelled vesicles (yellow vesicles+ white arrows). Scale bar = 5µm. Lower: Maximum projection by confocal imaging of live osteoblast (GFP) and GMP (CFP) co-culture demonstrating the transfer of PKH-26 labeled vesicles (yellow+white arrows) from osteoblasts to GMPs as indicated by the white arrows. Scale bar = 10µm. **(H)** Frequency of live progenitors labeled with PKH-26 vesicles from co-cultured osteoblasts. **(I, J)** FACS analysis of BM of Ocn-GFP mice injected with pHrodo, percentages are of parent gate **(I)** Granulocytic (Ly6G+) and monocytic (Ly6G-Ly6C+) cells gated on non-erythroid (CD71-Ter119-) BM **(J)** GMP^GFP+^, GMP^GFP-^. **(K)** Fold change in GFP+ cells post irradiation, 5FU and systemic *C. albicans* infection. Fold change is calculated from the mean of GFP+ cells frequency of two independent experiments as shown in fig. S2I-N. Data in **A, H** is presented as mean ± s.d. *\*\** p<0.01*, \*\*\** p*<0.001, \*\*\*\** p*<0.0001*.

Given that GMPs give rise to phagocytic cells (Akashi et al., 2000), we tested whether GMP^GFP+^ cells simply had greater phagocytic ability by injecting Ocn-GFP^Topaz^ mice with *E. coli* particles labeled with a pH-sensitive dye (pHrodo) that fluoresces within the acidic milieu of the phagosome (Lenzo et al., 2016). Phagocytic (pHrodo-positive) Ly6G^-^Ly6C^+^ monocytes and Ly6G^+^ granulocytes were GFP negative and hence were not labeled with EVs (Fig. 2I) while both GMP^GFP+^ and GMP^GFP-^ were not capable of phagocytosis (pHrodo-negative) (Fig. 2J). These data in combination with the data presented in Fig. 1 argue against the uptake of free unbound GFP by phagocytosis but rather through EV mediated transfer.

To further highlight the regulated nature of this process and rule out randomness, we tested the effect of three stress states on EV transfer to GMPs: genotoxicity from low dose *γ*-irradiation or 5-fluorouracil (5FU) and inflammation induced by systemic *C. albicans* infection. The frequency of GFP uptake was selectively increased in GMPs (1.5-2-fold) but not in CMPs or LKS 12 hours post exposure to the three stresses with no major changes in the absolute counts of GMPs (Fig. 2K, fig. S2I-Q). The increase in the frequency of GMP^GFP+^ cells was prior to the selective changes in the absolute numbers of total GMPs at 24 but not in CMPs or LKS (fig. S2R-T), consistent with an increase in EV uptake rather than rapid proliferation or differential BM retention of GMPs and highlighting a distinctive effect of EVs on GMPs under stress.

### tiRNAs are the most abundant sncRNAs in osteoblastic EVs

Extracellular vesicles carry proteins, lipids, metabolites and nucleic acids as cargo (Keerthikumar et al., 2016). The most abundant nucleic acids in EVs are mRNAs and small non-coding RNAs (sncRNA) (Valadi et al., 2007; Wei et al., 2017). The sncRNA content of BM-derived EVs and of GMP^GFP+^ and GMP^GFP-^ from the Ocn-GFP^Topaz^ mouse model was analyzed by RNA sequencing (Fig. 3A). The vast majority (85% of reads) of EV sncRNA consisted of tRNAs (Fig. 3B, table S2), with tRNAs coding for Gly-GCC-2, Glu-CTC-1 and Gly-CCC-5 as the most abundant (Fig. 3C, fig. S6, table S1). Among EV miRNAs, mir-148, let-7i and mir-143 were the most represented (fig. S3A).

**Fig. 3:**
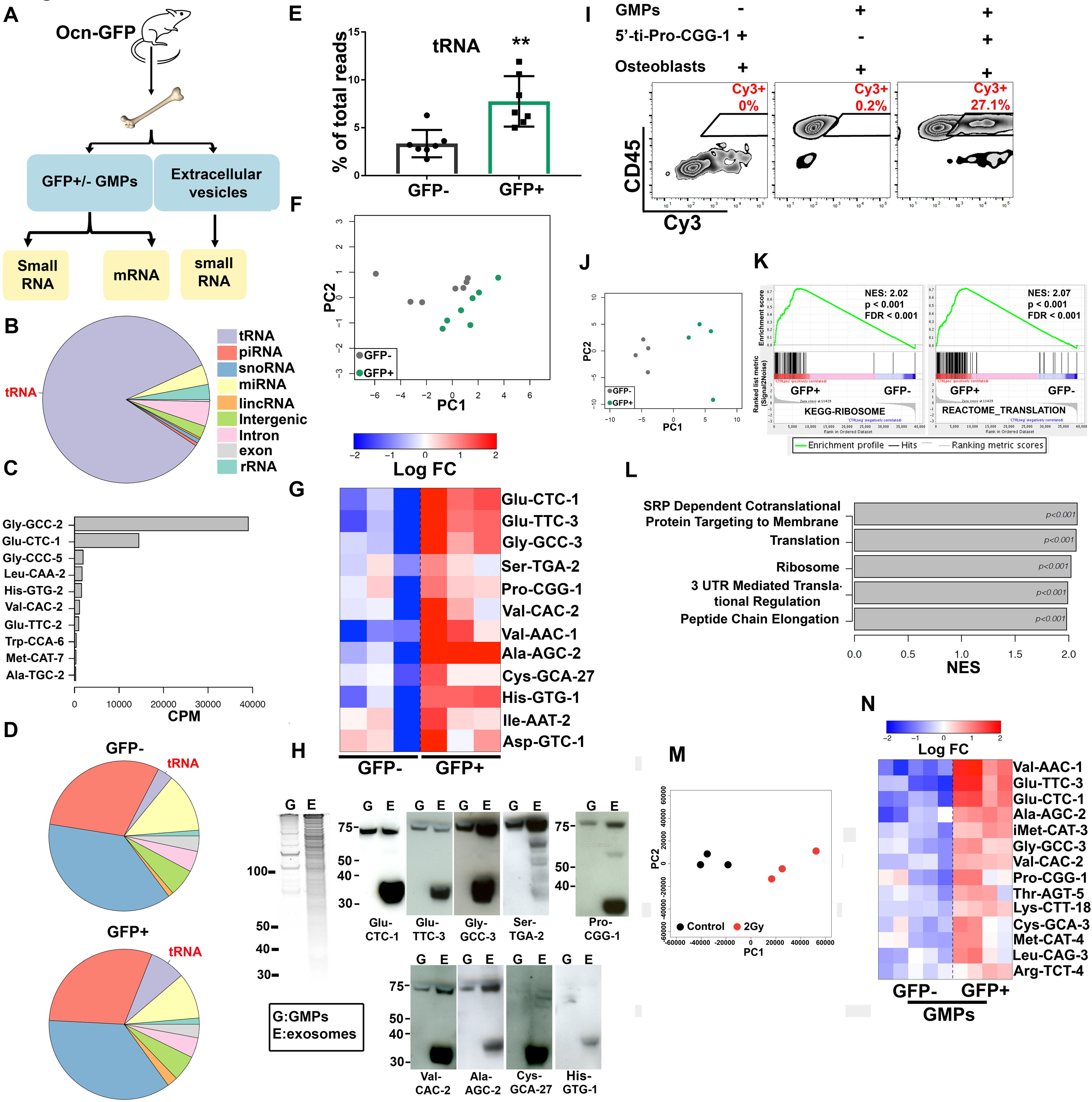
tiRNAs are the most enriched small RNAs in murine BM derived EVs. (A) Overview of RNA sequencing experiment using Ocn-GFP animals. (B) Fractions of small RNA sequencing reads mapped to genomic elements in BM EVs. (C) Top ten tRNAs ranked by their abundance in BM EVs. Data represents three biological replicates. (D) Fractions of small RNA sequencing reads mapped to genomic elements in GMP^GFP+^ and GMP^GFP-^. Data represent 7 biological replicates. (E) Percent of reads mapping to tRNAs in GMP^GFP+^ and GMP^GFP-^. Data is presented as mean ± s.d. ** p<0.01. (F) Principal component analysis (PCA) based on tRNAs expression in GMP^GFP+^ and _GMP_GFP-. (G) Heatmap of tRNAs that are more abundant in GMP^GFP+^, > 1.5 fold change. The levels are shown as relative to the average abundance of a given tRNA across all samples. Given extremely high sequence similarity between tRNA species sharing the same anticodon (Extended Data Fig. 6, Table 1, column “DNA sequence”), we used one individual tRNA representative per group. Data represents one of two independent experiments. (H) Left, Sybr gold stained RNA gel, 750ng total RNA per sample. Right, northern blot analysis of small RNAs collected from total GMPs (G) and BM EVs (E). (I) Transfer of synthetic Cy3 labeled 5’-ti-Pro-CGG-1 from transfected primary osteoblasts to co-cultured GMPs. (J) PCA of transcriptome-wide gene expression levels in GMP^GFP+^ and GMP^GFP-^, based on mRNA sequencing. (K) GSEA enrichment plots for ribosomal and translation-related genes. (L) Top gene sets enriched in GFP+ cells according to GSEA. (M) Principal component analysis (PCA) based on tRNAs expression in control and irradiated BM EVs . (N) Heatmap of tRNAs with > 1.5 fold change comparing GMP^GFP+^ to GMP^GFP-^, in 2Gy irradiated Ocn-GFP mice. The levels are shown as relative to the average abundance of a given tRNA across all samples. Given extremely high sequence similarity between tRNA species sharing the same anticodon (Extended Data Fig. 6, Table 1, column “DNA sequence”), we used one individual tRNA representative per group.

In GMP^GFP+^ and GMP^GFP-^, the majority of sncRNA were piRNAs and snoRNAs while miRNAs were more abundant than tRNAs (Fig. 3D, table S2). This finding is similar to published reports of cultured human mesenchymal cells (Baglio et al., 2015), glioma cells (Wei et al., 2017), T cells (Chiou et al., 2018), HEK293T (Shurtleff et al., 2017). Interestingly, the overall level of tRNAs was more than two-fold higher in GMP^GFP+^ compared to GMP^GFP-^, a distinctive finding among the sncRNA (Fig. 3E, fig. S3C, table S2). In addition, the GMP^GFP+^ and GMP^GFP-^ cells had distinct tRNA species levels by principal component analysis (PCA) (Fig. 3F). Twelve tRNAs had significantly higher levels in GMP^GFP+^ compared to GMP^GFP-^ in two independent experiments (Fig. 3G, fig. S3D) and were detected in BM EVs (table S1). EVs derived from cultured primary osteoblasts were also dominated by tRNA (90% of small RNA reads) and were markedly increased compared to tRNAs in the originating osteoblasts (fig. S3B, table S2). Val-AAC-1, Ser-TGA-2, Pro-CGG-1, Glu-TTC-3, Glu-CTC-1 and His-GTG-1 were particularly abundant in osteoblast derived EVs (fig. S3E, table S1.

Northern blot (NB) analysis on total RNA from BM derived EVs confirmed the presence of seven out of the top ten differentially abundant tRNAs within EV-labeled GMPs (Fig. 3H). Interestingly, smaller tRNAs of around 35 nucleotides were much more abundant than certain mature tRNAs within BM EVs and could not be detected in CD45+ or CD45-BM cellular RNA (Fig. 3H; fig. S3F). These smaller tRNAs had the size of tiRNAs, originally considered a byproduct of tRNA degradation (Borek et al., 1977; Speer et al., 1979), but increasingly recognized as a regulated tRNA processing product modulating protein translation (Anderson and Ivanov, 2014; Fricker et al., 2019; Kim et al., 2017; Yamasaki et al., 2009). Through their effect on translation, tiRNAs enable cell tolerance of stress conditions including oxidation, UV irradiation, heat shock and starvation (Fricker et al., 2019; Ivanov et al., 2011; Yamasaki et al., 2009). Probes for Cys-GCA-27, His-GTG-1 detected only tiRNA (not tRNA) within EVs (Fig. 3H).

We confirmed the transfer of tiRNAs from osteoblasts to GMPs through a co-culture assay of primary GMPs and primary osteoblasts producing Cy3 labeled synthetic 5’ tiRNA Pro-CGG-1 (5’ti-Pro-CGG-1) (Fig. 3I). Together, these findings are consistent with EV transfer of select tiRNAs from osteoblastic cells to GMPs.

To investigate the effect of the transferred small RNAs on the recipient GMPs, we performed mRNA sequencing of GMP^GFP+^ and GMP^GFP-^, which revealed distinctly different patterns of gene expression, as shown by PCA (Fig. 3J), with 21 significantly upregulated and 108 downregulated mRNAs (fig. S4, table S3). Pathway enrichment analysis using GSEA(Subramanian et al., 2005) indicated the upregulation of ribosomal and protein translation related genes in GMP^GFP+^ cells (Fig. 3K-L). Sequencing of EV mRNA was not performed due to diminished ribosomal RNA peaks, a finding that has been reported by others (Wei et al., 2017). To investigate the effect of stress on the tRNA content of EVs and GMP^GFP+^, small RNA sequencing was performed on BM EVs and GMPs from Ocn-GFP animals 12 hours post irradiation (2Gy). Principal component analysis and individual gene expression levels demonstrated a distinct tRNA content between EVs from control and irradiated animals (Fig. 3M, table S1). Analysis of the GMPs detected 14 tRNAs that are significantly more abundant in irradiated GMP^GFP+^ compared to GMP^GFP-^ from the same animals (Fig. 3N).

### Osteoblastic EVs tiRNA cargo enhances protein translation and proliferation in recipient GMPs

To validate this upregulation of protein synthesis machinery, we performed an *in vivo* protein translation assay by injecting Ocn-GFP^Topaz^ mice with O-Propargyl-puromycin (OPP), a molecule incorporated into nascent peptides that enables flow cytometric measurement of protein synthesis rates (Liu et al., 2012). In agreement with our pathway analyses, a significant increase in protein synthesis was observed in GMP^GFP+^ cells (Fig. 4A-B). These findings have two potential explanations: (1) cells with high protein synthesis preferentially take up EVs or (2) EV uptake leads to higher protein translation. To discriminate between these, we used a model in which the expression of *Homeobox-A9* (*HoxA9*) results in the differentiation arrest of primary mouse GMPs at a self-renewing stage enabling clones of a uniform cell stage and phenotype to be isolated and expanded (Sykes et al., 2016). This system enables a uniform population of GMP to be adoptively transferred and the *in vivo* consequences of EV content transfer evaluated.

**Fig. 4:**
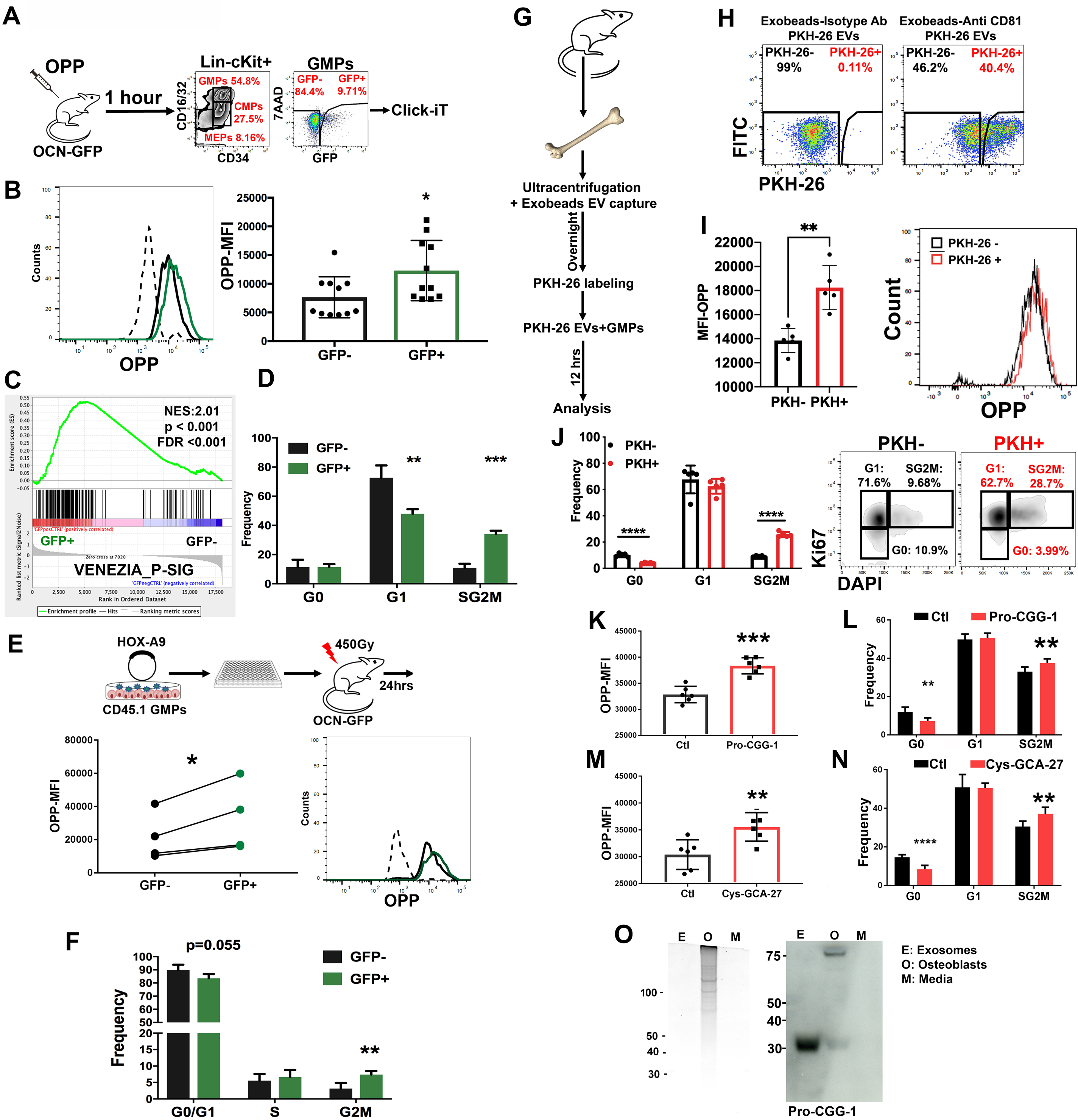
Osteoblastic cell derived EVs induce cell proliferation and increased protein translation in recipient GMPs through the transfer of tiRNAs. **A-D** Analysis of EV-labeled GMPs (GMP^GFP+^) for **(A, B)** the incorporation of OPP. Data represents two independent experiments **(C)** GSEA analysis and **(D)** cell cycle analysis, n=3. Data represent one of two independent experiments. **E, F** analysis of clonally derived myeloid cell line **(E)** OPP incorporation, analysis was performed using a paired student t-test and **(F)** cell cycle analysis, n=4. Data represent one of 2 independent experiments. **(G,H)** uptake of PKH-26 labeled BM EVs by live GMPs in culture. **(I)** enhanced OPP incorporation and **(J)** cellular proliferation in GMPs that take up PKH-26 labeled EVs. **(K, M)** OPP incorporation and **(L, N)** cell cycle analysis of synthetic Cy3 labeled tiRNA or control piRNA transfected GMPs. Analysis is performed on live, Cy3+ cells, n=6. Data represent two independent experiments. (O) Left, Sybr gold stained RNA gel, 75ng total RNA for EVs (E) and media (M) samples and 2ug for the Osteoblast (O) sample. Right, northern blot analysis of small RNAs collected from osteoblasts and their EVs released in the culture media. Data in **B,D,F,I-N** is presented as mean ± s.d. * p<0.05, ** p<0.01, *** p<0.001, **** p<0.0001.

Sub-lethally irradiated Ocn-GFP^Topaz^ mice were transplanted with clonal CD45.1-*HoxA9* GMP progenitors. One day post-transplantation, GFP was detected in the adoptively transferred cells. Further, GFP+ cells had a significantly higher rate of protein translation (by OPP analysis) compared with GFP-cells. These data with uniform starting GMPs, suggest that protein translation is directly induced by the transfer of EV-contents and argue against intrinsic differences in cells leading to selective uptake (Fig. 4E).

We hypothesized that the increased rate of protein translation would correlate with an increased rate of cell cycling. Indeed, a molecular signature of proliferating hematopoietic stem cells (Venezia et al., 2004) was enriched in GMP^GFP+^ by GSEA analysis (Fig. 4C). The GMP^GFP+^ demonstrated an increased frequency of cells in the S/G2M phase of cell cycle (>3-fold increase), as indicated by Ki67-staining (Fig. 4D and fig. S5A). The GFP+ clonal *HoxA9* GMPs also had increased cell cycling *in vivo* (Fig. 4F and fig. S5B). To further confirm the specificity of this phenotype to EV uptake, we isolated BM EVs by ultracentrifugation followed by anti-CD81 magnetic bead capture and added them to cultured GMPs 12 hours before analyzing for protein translation and cell cycle. Analysis by flow cytometry confirmed the uptake of EVs captured by anti-CD81 coated beads but not by an isotype control (Fig. 4G-H). Cells labeled by the EVs demonstrated an enhanced rate of protein translation (Fig. 4I) and cellular proliferation (Fig. 4J). Since tiRNAs are enriched in mouse BM EVs, we tested whether the tiRNA equivalents of the top ten differentially abundant tRNAs in GMP^GFP+^ increased protein translation and cell cycling. Synthetic tiRNAs or a piRNA control sequence (5’-phosphorylated and 3’-Cy3-labeled) were transfected into primary mouse GMPs; protein translation and cell cycle were assessed 24-hours post-transfection. 5’-ti-Pro-CGG-1 and 5’-ti-Cys-GCA-27 significantly increased the rate of protein translation in Cy3+ cells while the other tiRNAs did not (Fig. 4K, M, fig. S5C, E). Similarly, 5’-ti-Pro-CGG-1 and 5’-ti-Cys-GCA-27 increased the frequency of cells in the S/G2M phase of the cell cycle while the other tiRNAs did not except for 5’-ti-His-GTG-1 which decreased the frequency of cells in the S/G2M phase and increased those in G0. However, given its low abundance in BM EVs we believe its effect is minor compared to 5’-ti-Pro-CGG-1 and 5’-ti-Cys-GCA-27 that are much more abundant (Fig. 4L, N, Fig. 3H and fig. S5D, F). Notably, 5’-ti-Pro-CGG-1 was present in EVs isolated from primary osteoblasts by NB, whereas the mature tRNA Pro-CGG-1 (m-Pro-CGG-1) was not. In osteoblast cellular RNA, both m-Pro-CGG-1 tRNA and 5’-ti-Pro-CGG-1 were detected; however, the tiRNA was significantly less abundant in the cells than in the EVs (Fig. 4O). In contrast, neither 5’-ti-Cys-GCA-27 nor m-Cys-GCA-27 were detected in primary osteoblast EVs (data not shown) indicating a non-osteoblastic source for the 5’-ti-Cys-GCA-27 detected in total BM EVs. These data indicate that m-Pro-CGG-1 might be processed in EVs or 5’-ti-Pro-CGG-1 is sorted into EVs and that it is the tiRNA fraction that likely drives changes in EV-recipient cells. Notably, Pro-CGG-1 was differentially abundant in GMP^GFP+^ compared to GMP^GFP-^ upon irradiation pointing towards its potential role in response to stress.

### Increased osteoblastic EVs enhance response to stress

In light of the increased osteoblast derived EV transfer to GMPs under stress followed by GMP expansion (Fig. 2K, fig. S2I-K, R-T), we tested whether 5’-ti-Pro-CGG-1 could affect GMP differentiation *in vitro* when compared to the piRNA control sequence and demonstrated an enhanced rate of differentiation by immune-phenotype (increased frequency of Ly6g+CXCR2+ granulocytic and CD11b+CX3CR1+ monocytic cells), functional phagocytosis of pHRodo labeled *E. coli* and *C. albicans* killing in differentiated cells (Fig. 5A-G, fig. S7A) in addition to the increased cell cycling and protein translation previously noted (Fig. 4K-N). These data provide a potential role for osteoblast derived EVs and their cargo in tuning GMP stress response in a regulated manner. To investigate this in an *in vivo* setting and since there are no robust methods to specifically inhibit or enhance osteoblastic EV transfer *in vivo*, we increased the number of sender osteoblastic cells and measured the effect on EV transfer and myeloid based immunity *in vivo*.

**Fig. 5:**
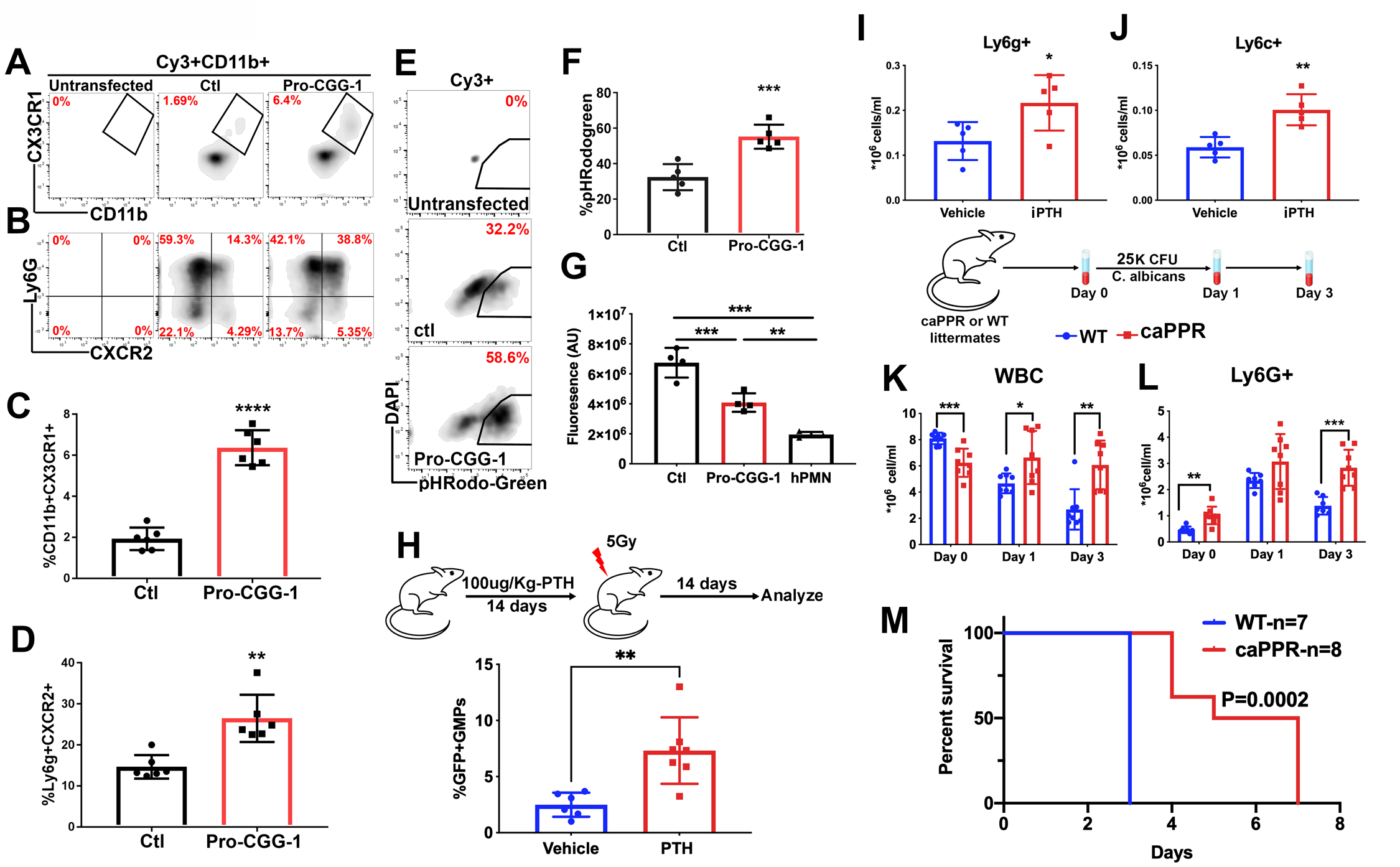
Enhanced response to stress in mice with increased osteoblast derived EVs. **(A-D)** phenotypic analysis by flow cytometry of 5’-ti-Pro-CGG-1 or piRNA control transfected GMPs, gates are on Cy3+, CD11b+ cells **(A, C)** monocytic **(B, D)** granulocytic markers. Data represent two independent experiments. **(E, F)** Phagocytosis assay analysis. Gates are on Cy3+ cells. Data represent two independent experiments. **(G)** Fluorescence signal from metabolically active *C. albicans* co-cultured with Cy3+ GMPs for two hours. Data represent one of two independent experiments. Analysis was performed using one-way ANOVA with no correction for multiple comparisons. **(H)** Frequency of GMP^GFP+^ (parent gate) after 14 days of iPTH injections. Data represent independent biological replicates of two independent experiments. **(I)** Peripheral blood neutrophils **(J)** monocytes 14 days post irradiation in iPTH treated mice. Data represent two independent experiments. **(K)** Peripheral blood WBC **(L)** and neutrophil counts in *C. albicans* infected caPPR mice. **(M)** survival analysis in caPPR mice post *C. albicans* infection. Data represent one of two independent experiments. Data in **C, D, F, G-L** is presented as mean ± s.d. *p<0.05, **p<0.01***p<0.001, ****p<0.0001.

This was achieved either pharmacologically using intermittent recombinant PTH (iPTH) injection (Silva et al., 2011) or genetically by using the osteoblast specific constitutively active PTH and PTH related peptide receptor (caPPR) mouse model under the control of the collagen 1 promoter (Calvi et al., 2001). Intermittent PTH injection increased osteoblasts, osteoblast derived EV transfer to GMPs and enhanced myeloid cell recovery two weeks post radiation injury as reflected by significantly higher neutrophils and monocytes (Fig. 5H-J, fig. S7B,C). Since iPTH may directly affect hematopoietic cells among many others, we used the caPPR mice which similar to iPTH injection demonstrated increased osteoblasts as well as increased EV transfer to GMPs in mice crossed with the Ocn-GFP^Topaz^ reporter (fig. S7D,E). When challenged with a lethal dose of *C. albicans*, CaPPR mice demonstrated a sustained increase in myeloid cell response (Fig. 5K-L, fig. S7F-G) and, notably, improved survival (Fig. 5M, fig. S7H).

## Discussion

Our findings identify an unconventional mechanism through which mesenchymal cells in the BM regulate the highly dynamic myeloid component of innate immunity and identify tiRNAs as an EV cargo that can alter the physiology of recipient cells. We demonstrate in an *in vivo* setting through the use of reporter mice that label specific BMMS that osteoblastic cells within the BM communicate with hematopoietic progenitors via EVs, transmitting complex information through sncRNA. Exchange of EVs between mesenchymal cells and hematopoietic cells has been reported in a non-physiological *in vitro* settings or by transferring *in vitro* generated EVs to animals (Goloviznina et al., 2016; Morhayim et al., 2016; Wen et al., 2016a). However, here we show that the process occurs *in vivo* and is modulated by stress. Further, we provide *in vivo* as well as *in vitro* evidence that select mesenchymal cells have a higher ability to produce and transfer EVs with preferential uptake by specific hematopoietic progenitors. This does not rule out the existence of EV mediated communication between other BM stromal populations (endothelial, hematopoietic, nervous system) that are not targeted by the mesenchymal reporters used and HSPCs. The cargo of tiRNA results in vesicular signaling that alters fundamental behaviors such as cell cycle and protein translation. Specifically, 5’-ti-Pro-CGG-1 enriched in osteoblast derived EVs, can enhance protein translation, cellular proliferation and eventually differentiation in recipient GMPs. These phenotypic changes occur without the complex signal transmission and transcriptional regulation that are necessary downstream components of traditional ligand-receptor interactions. In this way, specific stromal cells provide a stress regulated means of directly transferring tiRNA to activate key programs of cell physiology. By enhancing protein translation, activating cell proliferation in specific myeloid progenitor cells, this tiRNA transfer augments defense against pathogens like the *Candida* tested here.

EVs with tRNA and tiRNA contents have been reported previously in cell lines but we show that this is occurring *in vivo* in a manner that modulates the organisms response to physiologic stress (Baglio et al., 2015; Chiou et al., 2018; Jeppesen et al., 2019; Shurtleff et al., 2017; Wei et al., 2017). We demonstrate that the extent of EV transfer can be modified *in vivo* by altering the producer cells. This resulted in improved myeloid response and infection control.

The impact of tiRNA on protein translation that we observed was not anticipated. 5’ tiRNAs with oligoguanine terminal motifs were initially shown to inhibit protein translation through the displacement of the cap binding complex eIF4F (Ivanov et al., 2011; Yamasaki et al., 2009).

However, evidence has now emerged that tRNA fragments can have highly pleiotropic effects (Fricker et al., 2019; Kim et al., 2017) that may be further modulated by post transcriptional modifications such as pseudouridylation (Guzzi et al., 2018) and methylation (Blanco et al., 2016; He et al., 2021; Schaefer et al., 2010; Tuorto et al., 2012). mRNA stability, Argonaute/PIWI and other RNA binding proteins and ribosome/mRNA interactions may all be affected by tiRNA such that the impact of tiRNAs is highly cell context dependent (Magee and Rigoutsos, 2020).

Extracellular vesicles bearing tiRNA add to the repertoire of mechanisms by which niche cells can modulate parenchymal cell responses to stress, providing a mechanism that is more direct and likely more immediate than cytokine-receptor interactions. We hypothesize that this sncRNA exchange may be more primitive evolutionarily and by virtue of its potential to exchange complex admixtures of regulatory RNAs, more nuanced in the effects it can induce. Furthermore, EV transfer of sncRNA may account for prior findings that the loss of small RNA processing enzymes in stromal cells can result in hematopoietic dysfunction (Goncalves et al., 2016; Raaijmakers et al., 2010). Non-coding RNA signaling is made possible by direct exchange of cell microparticles and represents a distinctive form of stress modulated communication between niche and parenchymal cells that affects normal and aberrant tissues and may change organismal physiology to challenges such as infection.

### Limitations of study

We realize that our methods to manipulate the levels of EVs are indirect and therefore the results should be interpreted with caution. Molecules or genetic models that could specifically abrogate or increase EV release *in vivo* in a specific population of cells without a wider effect on essential signaling pathways remain to be identified. Without tools to experimentally manipulate EV release directly, our indirect method is consistent with EVs inducing the effect, but not definitive of it. Furthermore, we cannot rule out the possibility that EV recipient cells upregulate tRNA processing into tiRNAs due to the lack of a comprehensive list of the enzymes that drive this process. Finally, it would be exciting to identify the elements responsible for the high level of EV release by osteoblasts in comparison to other mesenchymal subsets and the tropism of these EVs to GMPs. Such knowledge would strengthen the translational impact of our findings potentially allowing for engineering EVs that could be used as delivery method for gene editing machineries.

## Acknowledgements

The authors would like to thank Phillip Chea at Massachusetts General hospital for his technical help; Adalis Maisonet and Ulandt Kim from the MGH next generation sequencing core (NIH P30 DK040561) for helping in library constructions and Illumina sequencing; Scott Mordecai at the department of pathology flow, image and mass cytometry core (1S10OD012027-01A1) at MGH for his help in imaging flow cytometry. We would also like to acknowledge the HSCI-CRM flow cytometry facility for their help in flow cytometry analysis and sorting; Center for Skeletal Research Core (NIH P30 AR066261) for their help in microcomputed tomography; Microscopy Core of the Center for Systems Biology/Program in Membrane Biology, (DK043351) and (BADERC award DK057521).

## Author contributions

Y.K. initiated the project, designed and performed experiments, and wrote the manuscript. F.J., A.A., R.I.S. performed bioinformatics analyses. M.M. and Y.A. performed the northern blots.

D.B.S. helped establish the HA9 cell line and provided thoughtful advice. A.P., J.S, T.B, F.M., N.S., K.K., A.K.S. and B.S. helped perform experiments. M.K.M and P.I. provided thoughtful advice. D.T.S. initiated the project, provided supervision and wrote the manuscript. All authors read and helped edit the manuscript.

## Data and materials availability

The datasets generated during the current study and analyzed in Fig. 3, fig. S3, fig. S4, Fig. S6 have been submitted to Geo (GSE127872).

## Methods

### Mice

All animal experiments were approved by the Institutional Animal Care and Use committee at Massachusetts General Hospital. Wild type CD45.2 (C57BL/6J), congenic CD45.1 (B6.SJL-Ptprc<A >Pepc /BoyJ), CAG-ECFP (B6.129(ICR)-Tg(CAG-ECFP)CK6Nagy/J) and Rosa26- YFP (Rosa-YFP, B6.129X1-*Gt(ROSA)26Sor^tm1(EYFP)Cos^*/J) mice were purchased from Jackson Laboratories. Col2.3-GFP(Kalajzic et al., 2003), Ocn-GFP^Topaz^ (Bilic-Curcic et al., 2005), Nes-GFP(Mignone et al., 2004), Osx-Cre::GFP(Rodda and McMahon, 2006), caPPR (Calvi et al., 2001) and Oc-Cre(Zhang et al., 2002) were previously described. Gender matched mice, 10-14 weeks of age were used in all experiments unless stated otherwise.

For total BM transplant experiments, mice received 2X(6.5Gy) doses from a cesium-137 irradiator within a 4 hours period. The day after, 1x 10^6^ BM nucleated cells were transplanted via retro-orbital injection. Mice were analyzed 8 weeks post-transplantation. For the clonal cell line transplant, mice received a dose of (4.5Gy). The day after, the mice received 2 X (20*10^6^) cells 8 hours apart and mice were analyzed one day after.

For genotoxic stress, mice received a dose of (2Gy or 5Gy) or one intraperitoneal injection of 150mg/Kg 5FU. For systemic fungal infection, (C57BL/6J mice received 100K CFU and CaPPR mice received 25K of *C. albicans* (SC5314) in 200ul PBS through the tail vein. Mice were analyzed 12 hrs later.

For iPTH injection, mice were given 14 daily subcutaneous injections of vehicle (10mM citric acid, 150nM NaCl, 0.05% Tween 80) or 100ug/Kg body weight of Y^34^hPTH(1-34) amide (SVSEIQLMHNLGKHLNSMERVEWLRKKLQDVHNY.NH2)

### Genotyping and QPCR

Mouse tail DNA was used for genotyping using the indicated primers: For Ocn-GFP^Topaz^, Col1-GFP and Nes-GFP: F: 5’ CTG GTC GAG CTG GAC GGC GAC GTA AC 3’ R: 5’ ATT GAT CGC GCT TCT CGT TGG GG 3’

For Osx-GFP: F: 5’ CTC TTC ATG AGG AGG ACC CT 3’ R: 5’ GCC AGG CAG GTG CCT GGA CAT 3’ For R26-YFP:

R26-YFP-WT: 5’ GGAGCGGGAGAAATGGATATG 3’

R26-YFP-Common: 5’ AAAGTCGCTCTGAGTTGTTAT 3’

R26-YFP-Mutant: 5’ AAGACCGCGAAGAGTTTGTC 3’

For Oc-Cre:

Mut-F: 5’ GAC CAG GTT CGT TCA CTC ATG G 3’ Mut-R: 5’ AGG CTA AGT GCC TTC TCT CTA CAC 3’

For CaPPR:

Col1: 5’ GAGTCTACATGTCTAGGGTCTA 3’

G2: 5’ TAGTTGGCCCACGTCCTGT 3’

### Flow cytometry analysis and sorting

Mice were sacrificed through CO2 asphyxia. Whole BM mononuclear cells (MNCs) were collected by crushing of bones (tibias, femurs, hips, humeri and spine) and subjecting the cells to density gradient centrifugation (Ficoll-Paque Plus, GE Healthcare) at 400g for 25 minutes with brakes turned off. Mononuclear cells were then stained in PBS supplemented with 2%FBS using the following antibodies: CD45-APCCy7 (30F-11), Sca1-BV421 (D7), cKit-BuV395 or APCCy7 (2B8), CD16/32-BV605 or PeCy7 (2.4G2), CD34-AF647, Pe or FITC (RAM34), IL7R-Pe (A7R34), Biotinylated lineage cocktail (CD8A (53-6.7), CD3E (145-2C11), CD45R (RA3-6B2), GR1 (RB6-8C5), CD11b (M1/70), Ter119 (Ter-119), CD4 (GK1.5) followed by Streptavidin- BV711 conjugate. CD45.1-BV650 (A20) was used for chimerism in transplant experiments. To assess EV transfer in the mature compartment of the BM, total BM cells were stained using Ter- 119-Pe (Ter-119), CD71-Pe (R17217), CD11b-AF700 (M1/70), CD3e-APC (145-2C11), CD45R-eFluor450 (RA3-6B2) 7-Aminoactinomycin D (7AAD) was used as a viability dye. At least 2x10^6^ events were collected per sample for stem and progenitor cell analysis using a BD FACSARIA I, II or II for both analysis and sorting. Analysis was performed using the FlowJo software.

For bone analysis by flow, bones (tibias, femurs, hips, humeri and spine) were cut into small pieces after crushing and digested for one hour at 37^ο^C in a shaking water bath at 120rpm. The flow through was strained over 70um strainer, washed and stained with antibodies for Ter119-PeCy7 9Ter119), CD45-peCy7 (30F-11), CD31-APC (MEC 13.3).

### Extracellular vesicle collection

For RNA extraction from EVs, mice were euthanized, and BM was flushed in PBS from tibias, femurs, hips and humeri. For the collection of cultured osteoblast EVs, 500K YFP+ osteoblasts were cultured in *α*-MEM supplemented with 10% FBS, 1% Penicillin/Streptomycin, 1% L-Glutamine 50ug/ml ascorbic acid (Sigma) until cells reached 80% confluency. Media was removed and cells were washed twice with pre-warmed PBS. Fresh *α*-MEM supplemented with 2% exosome free FBS, 1% Penicillin/Streptomycin, 1% L-Glutamine, 50ug/ml ascorbic acid (Sigma) was added for three days after which media was collected for EV isolation. Cells were excluded by centrifugation for 5 minutes at 500g. EVs and RNA were then isolated from the supernatant using the Exoeasy and miRneasy (Qiagen) according to manufacturer’s instructions. For nanoparticle tracking analysis (NTA), electron microscopy and WB after cell exclusion, the supernatant was transferred into a new tube and centrifuged for 20 minutes at 20,000g. The supernatant was then passed through a 0.22µm low protein binding filter and subjected to ultracentrifugation at 120,000g using the SW32Ti rotor using the Optima L90K ultra-centrifuge from Beckman coulter for 120 minutes. Pellets were washed once with PBS followed by a second round of ultracentrifugation.

For culture with primary GMPs, protein quantification was performed using the DC protein assay (Biorad). 100ug were added to 50K GMPs sorted the day before and cultured in StemSpan SFEMII supplemented with 1% L-Glutamine and Penicillin/Streptomycin with no cytokines (Stem cell technologies). Cells were cultured in a humidified incubator at 37°C and 5% CO2 for 12 hours and then washed twice with PBS-2%FBS with 7AAD. Live cells were sorted using a BD FACS ARIA II.

### Nanoparticle tracking analysis (NTA)

Following PBS wash and ultracentrifugation, EV pellets were analyzed using Nanosight instrument technology (NTA 3.2 Dev Build software) (5X60 seconds video/sample, detection threshold: 5) for nanparticle size.

### Confocal microscopy

GFP+/- LKS and GMPs were sorted as described above and live cells were imaged in 8 chamber borosilicate coverglass system (nunc) coated with human plasma fibronectin (EMD Millipore) using a Leica TCS SP8 confocal microscope equipped with two photomultiplier tubes, three HyD detectors and three laser lines (405nm blue diode, argon and white-light laser) using a 63x objective at 200Hz and bidirectional mode. 8-bit images were acquired at 512x5212 resolution and processed by Imaris software (Bitplane). For co-culture, 25*10^3^ PKH-26 labeled primary osteoblasts / were cultured in 8 chamber borosilicate coverglass system (nunc). Sorted GMPs from Actin-CFP mice were co-cultured overnight before imaging.

### EVs Exobead capture and PKH-26 labeling

Extracellular vesicles were prepared by ultracentrifugation as described above and washed once with PBS. EVs were then pulled down by incubating with anti CD81-Biotin (Eat-2, Biolegend) coated streptavidin beads overnight rotating at 4°C (Exobeads, Invitrogen). Beads were then collected using a magnet and washed 3 times with PBS supplemented with 0.1% BSA. For fluorescent labeling, pulled down EV/Bead complexes were stained using anti CD9-AF647 (MZ3- Biolegend) and analyzed using BDFACS ARIA II. For PKH-26 (Sigma-Aldrich) labeling, 200ug of ultracentrifugation enriched EVs were pulled down using anti-CD81 coated Exobeads as described above. Captured EVs were labeled in 200ul volume for five minutes. Labeling was stopped using an equal volume of PBS with 1% BSA and samples were washed three times according to manufacturer’s instructions. The equivalent of 100ug starting material of Exobead captured EVs labeled with PKH-26 were added to 50K sorted GMPs in StemSpan supplemented with 1% Penicillin/Streptomycin and L-Glutamine without cytokines. Cells were analyzed 12 hours later for protein translation and cellular proliferation.

### Colony forming assay

Equal numbers of cells were sorted as described above and reconstituted in MethoCult (M3434- Stem Cell Technologies) according to manufacturer’s instructions or (M3234-Stem Cell Technologies) supplemented with 2ng/ml mIL3 and mIL6, 10ng/ml mSCF, 1U/ml hEPO. Recombinant cytokines were purchased from PeproTech. Colonies were manually enumerated 10 days post seeding. Colony size was measured for at least 10 colonies in each biological replicate using ImageJ.

### Cytospins and Wright Giemsa staining

GMP^GFP+^ and GMP^GFP-^ were sorted as described above and 20K cells were immobilized on slides using the cytospin for 1 minute at 1000 rpms (Thermo Scientific Shandon) and were allowed to air dry. Slides were stained in 100% Wright-Giemsa (Siemens) for 2 min, and in 20% Wright-Giemsa diluted in buffer for 12 min. Stained cells were rinsed in deionized water, and coverslips were affixed with Permount prior to microscopy.

### Imaging flow cytometry

GFP+/- LKS were sorted from Ocn-GFP^Topaz^ as described above and then analyzed using Amnis ImageStream, EMD Millipore).

### Isolation of DNA and RNA

DNA for genotyping was isolated from tails or cells using DNeasy blood and tissue kit (Qiagen). RNA was isolated using the miRNeasy micro or RNeasy micro kits depending on the downstream application. All extractions were performed according to manufacturer’s instructions.

### Immunoblotting

Total BM EVs or nucleated cells were lysed in NuPAGE LDS lysis buffer (Life Technologies) and proteins were quantified using the DC protein assay (Biorad). 20ug total proteins were loaded per lane. Immunoblotting was performed using rabbit polyclonal anti-GFP (ab290-abcam) and rabbit monoclonal anti-TSG101 (EPR7130B-abcam).

### Transmission electron microscopy

#### Negative staining

EV suspensions were fixed in 2% paraformaldehyde and 10µl aliquots applied onto formvar-carbon coated gold mesh grids; specimens were allowed to adsorb for 10-20 minutes. Grids were contrast-stained in droplets of chilled tylose/uranyl acetate (10-15min) or in 2% aqueous phosphotungstic acid (30-90sec). Preparations were allowed to air-dry prior to examining in a JEOL JEM 1011 transmission electron microscope at 80 kV. Images were collected using an AMT digital camera and imaging system with proprietary image capture software (Advanced Microscopy Techniques, Danvers, MA).

### Immunogold staining

Following adsorption of 10µl aliquots of EV suspensions, grid preparations were either placed immediately on drops of primary antibody anti-TSG101, Abcam (EPR7130B), or anti-GFP (ab290-abcam) in DAKO antibody diluent). In case of GFP labeling, EVs were pre-treated briefly with filtered permeabilization solution (PBS/BSA/saponin) prior to incubation in primary antibody. Incubation in primary antibody occurred for at least 1 hour at room temperature. Grids were then rinsed on droplets of PBS and incubated in goat anti-rabbit IgG gold conjugate (Ted Pella #15727, 15nm) or (Ted Pella #15726, 10nm) at least 1 hour at room temperature. Grids were then rinsed on droplets of PBS, then distilled water, followed by contrast- staining for 10 minutes in chilled tylose/uranyl acetate. Preparations were air-dried prior to examining in a JEOL JEM 1011 transmission electron microscope at 80 kV. Images were collected using an AMT digital camera and imaging system with proprietary image capture software (Advanced Microscopy Techniques, Danvers, MA).

### mRNA and small RNA sequencing and analysis

RNA-seq libraries for gene expression were constructed using Clontech SMARTer v.3 kit (Takara). Small RNA libraries were constructed using NEBNext Multiplex Small RNA Library Prep Set for Illumina (New England Biolabs). mRNA and small RNA libraries were sequenced on Illumina HiSeq2500 instrument, resulting in approximately 30 million reads and 15 million reads per sample on average, respectively.

mRNA sequencing reads were mapped with STAR aligner(Dobin et al., 2013) using the Ensembl annotation of mm10 reference genome. Read counts for each transcript were quantified by HTseq(Anders et al., 2015), followed by estimation of expression values and detection of differential expressed using edgeR(Robinson et al., 2010) after normalizing read counts and including only those genes with CPM > 1 for one or more samples. Differentially expressed genes were defined based on the criteria of >2-fold change in expression value and false discovery rate (FDR) < 0.001. RPKM expression values were submitted to the GSEA tool(Subramanian et al., 2005) to analyze the enrichment of functional gene categories among differentially expressed genes.

To analyze small RNA data, adaptor trimming was performed by Trimmomatic(Bolger et al., 2014) and the reads at least 18 bp long were kept for further analyses, resulting in approximately million reads per sample on average. Sequencing reads were aligned to mm10 reference genome using BWA aligner. To quantify the expression of various RNA species, we used the Ensembl Mus_musculus GRCm38.87(Zerbino et al., 2018) annotation of lincRNAs, miRNAs, snoRNAs, and mRNAs; the annotation of tRNAs from GtRNAdb^43^; and the annotation of piRNAs from piRNABank(Sai Lakshmi and Agrawal, 2008). To identify differentially expressed miRNAs and tRNAs, their expression levels were quantified by miRExpress(Wang et al., 2015) and SALMON(Patro et al., 2017) respectively, followed by calling differentially expressed RNAs using edgeR(Robinson et al., 2010).

60 out of all 471 murine tRNA sequences annotated in GtRNAdb database(Chan and Lowe, 2016) were identified as differentially expressed between GFP- and GFP+ based on the criteria of >1.5 fold change in both batches (n=3 and n=4 respectively). To assess the pattern of coverage by mapped sequencing reads for individual differentially expressed tRNAs, we used the BWA mapper with default settings(Li and Durbin, 2009) to provide exact genomic locations of mapped reads, as the exact read mapping is not provided, by design, by the SALMON method used for the quantitation of gene expression. These patterns of coverage revealed that the majority of the small RNA reads covered 5’ regions of tRNA sequences (Extended Data Fig.6). Because tRNAs with the same anticodon sequence share extremely high sequence similarity, it was challenging to distinguish between expression levels of individual tRNAs within these groups. Among differentially expressed tRNAs, the individual members of groups with the same anticodon had sequence identity above 85%, consistent with our clustering by the CD-HIT(Fu et al., 2012) tool (Extended Data Table 1, column “DNA Sequence”).

Therefore, in presenting expression values and differentially expressed tRNAs, as well as in follow-up experiments, we used one individual tRNA representative per group to represent the whole group of similar tRNA species. Extended Data Fig. 6 shows the density of sequencing reads over the length of tRNA sequences for these tRNA groups in all experimental conditions. One representative sequence is shown for each group; the whole groups are indicated in Extended Data Table 1.

#### *In vivo* phagocytosis assay

Ocn-GFP^Topaz^ mice were injected intravenously with 50mg/kg of pHrodo labeled *E-Coli* particles (Invitrogen) and one-hour post injection mice were sacrificed, and BM MNCs were collected, stained and analyzed as described above.

#### *HoxA9* clonal cell line

The **MSCVneo-*HoxA9*** ecotropic retrovirus donated by Dr. David Sykes. The vector has been previously described(Calvo et al., 2000). GMPS were sorted as described above from CD45.1 and cells were cultured in a 12 well plate pre-coated with human fibronectin (EMD Millipore) in RPMI1640 media + 10% Fetal Bovine Serum (FBS), 1% Penicillin/Streptomycin, 1% L-Glutamine, 10ng/ml SCF, 5ng/ml IL-3, 5ng/ml IL-6. Cells were transduced 24 hours later with

**MSCVneo-*HoxA9*** retrovirus in the presence of 8ug/ml Polybrene. The transduction was performed by spinfection (1000g for 60 minutes at room temperature). Following the spinfection, the cells were maintained in media described above and 24 hours later, they were selected for 4 days with G418 (Geneticin, 1mg/ml) (Invitrogen) and later maintained in cytokine media with no selection. Two weeks post transduction, cells were sorted as single cells in 96 well plate and maintained in the cytokine supplemented media for 2 weeks. Wells containing colonies were expanded and one was used for the clonal *HoxA9* cell line experiment. All through, cells were maintained in a humidified incubator at 37°C, 5% CO2. Cell line is available upon request from investigators.

### tiRNA transfection of GMPs

GMPs were sorted as described earlier from WT (C57Bl6/J) and 50K cells were cultured in 0.5mls of StemSpan^TM^SFEMII (Stem cell technologies) supplemented with 1% L-Glutamine and Penicillin/Streptomycin in addition to mouse recombinant cytokines: 10ng/ml SCF, 100ng/ml TPO, 5ng/ml IL3 and IL6 (PeproTech). Cells were transfected the day after with 0.5ul of a 100uM stock Cy3 labeled RNA oligos using Lipofectamine Stem (Invitrogen) at a ratio of 1:2 according to manufacturer’s protocol. RNA oligos were ordered from IDT with a phosphorylated 5’ end and Cy3 labeled 3’ end with the following sequences:

Pro-CGG-1-GGCUCGUUGGUCUAGGGGUAUGAUUCUCGCUUCG His-GTG-1-GCCGAGAUCGUAUAGUGGUUAGUACUCUGCAUUGU Cys-GCA-27-GCGGGUAUAGCUCAGGGGUAGAAUAUUUGACUG Ala-AGC-2-GGGGGUGUAGCUCAGUGGUAGAGCGCGUGCUUA Ser-TGA-2-GUAGUCGUGGCCGAGUGGUUAAGGCGAUGGACUUG Gly-GCC-3-GCAUUGGUGGUUCAGUGGUAGAAUUCUCGCCUGCC

Glu-CTC-1-UCCCUGGUGGUCUAGUGGUUAGGAUUCGGCGCUCU Glu-TTC-3-UCCCUGGUGGUCUAGUGGCUAGGAUUCGGCGCUUU Val-CAC-2-GUUUCCGUAGUGUAGUGGUUAUCACGUUCGCCUCA Control (piRNA)-UGUGAGUCACGUGAGGGCAGAAUCUGCUC

Half media change was performed 8 hours post transfection and cells were analyzed 24 hours post transfection.

### *In vitro* protein translation assay

Transfected cells were counted and 75K cells were incubated in a humidified 37°C incubator for 30 minutes in media containing 20uM O-Propargyl Puromycin (MedChem express). Cells were stained with the fixable LIVE/DEAD^TM^ yellow stain according to the manufacturer’s protocol followed by fixation using the Fixation/Permeabilization kit (BD Biosciences). After fixation, cells were washed with PBS supplemented with 3% BSA (Sigma)and then permeabilized using 1X perm/wash buffer (BD). Cells were stained for the OPP using the Click-iT Plus Alexa Fluor 647 Picolyl azide kit (Invitrogen) according to manufacturer’s protocol and analyzed using BD-FACS ARIA II.

### *In vivo* protein translation assay

Mice were injected intraperitoneally with 50mg/Kg OPP and sacrificed one hour later. BM MNCs were harvested as described earlier for myeloid progenitor cell surface staining. GMP^GFP+^ and GMP^GFP-^ or clonal *HoxA9* cells were sorted directly in the fixation buffer from the Fixation/Permeabilization kit (BD Biosciences). Cells were then washed with PBS supplemented with 3% BSA followed by the Click-iT reaction as described above. Analysis was done using BD- FACS ARIA II.

### Cell cycle analysis

For the tiRNA transfected GMPs, 75K cells were harvested and stained for viability using the fixable LIVE/DEAD far red stain (Invitrogen) according to manufacturer’s protocol followed by fixation and permeabilization using the Fixation/Permeabilization kit (BD Biosciences). Cells were then stained overnight at 4°C in 1X perm/wash buffer with FITC mouse Ki67 set (BD Pharmingen #556026). Cells were then washed with 1X perm wash buffer and re-suspended in PBS supplemented with 1ug/ml 4′,6-diamidino-2-phenylindole

(DAPI) (Invitrogen) and 100ug/ml RNAse A (Sigma)and incubated at room temperature for 15 minutes before analyzing by flow cytometry using BD FACS Aria II. Gates were drawn based on isotype control.

For uncultured cells, GMP^GFP+^ and GMP^GFP-^ or clonal *HoxA9* cells were directly sorted into fixation buffer and cell cycle staining was performed as described above.

### *In vitro* differentiation phenotypic analysis

Primary GMPs transfected with tiRNAs as described above were analyzed 3 days post transfection. Cells were harvested and washed once with PBS-2%FBS. Cells were then blocked for 5 minutes at room temperature using anti-mouse CD16/32 Fc block (1/50) (BD Biosciences). Cells were then incubated with the staining (Ly6g-APCCy7 (1A8), CXCR2-APC (SA044G4), CD11b-AF700 (M1/70), Ly6c-BV570 (HK1.4), CX3CR1-AF400 (SA011F11), cKit-BuV395 (2B8)) mix for 30 minutes at 4°C, washed and re-suspended in PBS-2%FBS containing DAPI (Invitrogen) for viability and analyzed using BD-FACS ARIA II. Analysis for differentiated cells was performed on live Cy3+ cells gated based on non-transfected cells.

### *In vitro* phagocytosis assay

Primary GMPs transfected with tiRNAs as described above were analyzed 3 days post transfection. 100*10^3^ cells were incubated with pHRodo green labeled *E.coli* (Invitrogen) at a ratio of 1:10 (cells:bacterial particles) for one hour at 37°C shaking. Cells were then collected, washed twice with PBS-2% FBS and re-suspended in DAPI containing buffer for viability and analyzed using BD-FACS ARIA II. Phagocytosis was assessed in live Cy3+ cells gated based on non-transfected cells. For differentiation and phagocytosis analysis, mTPO was not added to the media.

### Primary osteoblast culture

Primary osteoblasts were prepared as previously described with minor modifications(Panaroni et al., 2015). Tibias, femurs, hips and humeri were collected from Oc-Cre hemizygous, R26YFP homozygous mice. BM was flushed and bones were cut into small pieces that were digested in serum free αMEM containing 2mg/ml Collagenase type II (Worthington, Lakewood, NJ) for 2 hours at 37°C in a shaking water bath. Bone chips were washed with serum free α-MEM and resuspended in α-MEM supplemented with 10% FBS, 50ug/ml ascorbic acid (Sigma), 1% Penicillin/Streptomycin and 1% L-Glutamine. Cells were incubated at 37°C in a humidified 5% CO2 incubator for one week after which the media was changed. Two weeks post seeding, the bone chips and adherent cells were trypsinized and digested at 37°C in a shaking water bath for 30 minutes in serum free α-MEM supplemented with 2mg/ml Collagenase type II. Cells were then stained with CD31-APC (MEC13.3) and CD 45-Pacific Blue (30-F11) and GFP+ CD31- CD45- osteoblasts were sorted using BD FACS Aria II and a 100um nozzle.

For co-culture, sorted osteoblasts were seeded in 24 well plate (50K/well), 24 hours later, cells were transfected with 0.5ul of 100uM stock Cy3 labeled tiRNA using lipofectamine Stem (Invitrogen) at a 1:2 ratio. Media was changed 8 hours post transfection .

For PKH-26 (Sigma-Aldrich) labeling, osteoblasts were labeled according to manufacturer’s instructions and seeded in 8 chamber borosilicate coverglass system (nunc) at 25K/chamber.

One day later, media was changed to 125ul 2% FBS a-MEM before hematopoietic progenitors were added in an equal volume of 2%FBS IMDM. Twelve hours later, the co-culture was imaged by confocal microscopy.

For EV harvest, 500K sorted osteoblast were seeded in 100mm dishes and incubated in a humidified 5% CO2 incubator until cells reached 80% confluency. Media was then replaced with α-MEM supplemented with 2% exosome free FBS (Gibco), 50ug/ml ascorbic acid, 1% Penicillin/Streptomycin, 1% L-Glutamine. Media and osteoblasts were harvested 3 days later and EVs were collected using Exoeasy kit (Qiagen). Total RNA was extracted using miRNeasy micro (Qiagen).

### Northern blot

RNA was separated by size using 15% Novex TBE-Urea gels (ThermoFisher, EC6885). The RNA gel was incubated in 20ml 0.5X TBE with 1x SYBR Gold Nucleic Acid Gel Stain (Invitrogen, S11494) for 20 minutes and imaged using alpha imager HP.

The RNA was then transferred to positively charged nylon membranes with 0.45μm pores (Roche, 11209299001). RNA was crosslinked to the membrane using a UV Stratalinker 1800 (Stratagene). The blot was pre-hybridized with DIG Easy Hybridization Buffer (Roche, 11603558001) for 30 minutes at 40°C and then hybridized with DIG-5’-labelled probe overnight at 40°C in a rotation hybridization oven (Techne). Anti-sense-tiRNA DNA oligos were ordered from IDT and labelled with DIG using the DIG Oligonucleotide Tailing Kit (Roche, 03353583910). Sequences of the probes are: 5’-Ala-AGC-2- AAGCACGCGCTCTACCACTGAGCTACACCCCC,

5’-Cys-GCA-27- AGTCAAATATTCTACCCCTGAGCTATACCCGC,

5’-His-GTG-1- AATGCAGAGTACTAACCACTATACGATCTCGGC,

5’-Pro-CGG-1- AAGCGAGAATCATACCCCTAGACCAACGAGCC,

5’-Ser-TGA-2- AGTCCATCGCCTTAACCACTCGGCCACGACTAC,

5’-Val-CAC-2- AGGCGAACGTGATAACCACTACACTACGGAAAC,

5’-Glu CTC-1-CGCCGAATCCTAACCACTAGACCACCAGGGA,

5’-Glu-TTC-3-AGCGCCGAATCCTAGCCACTAGACCACCAGGGA,

5’-Gly-GCC-3- GAGAATTCTACCACTGAACCACCCATGC

Membranes were washed twice with 2x SSC containing 0.1% SDS at room temperature for 5 minutes, followed by one 5-minute wash with 1x SSC containing 0.1% SDS at 40°C. Next, membranes were blocked with 10 mL of 1x blocking solution diluted in 1x Maleic Acid Buffer (Roche, 115857262001) with 0.3% TWEEN 20 for 30 minutes at room temperature. One unit of Anti-Digoxigenin-AP Fab fragments (Roche, 11093274910) was added to the blocking solution and incubated for 30 minutes at room temperature. The membrane was washed twice with 1x Washing Buffer (Roche, 115857262001) for 15 minutes. Membranes were briefly equilibrated with 10 mL 1x Detection Buffer (Roche, 115857262001). To detect DIG-labeled probing, 1 mL of CPD-Star (Roche, 12041677001) diluted 1:5 with 1x Detection Buffer was applied to the membrane and exposed to autoradiography film (Amersham, 28906845) in the dark.

### C. albicans culture

Candida albicans, wild-type strain SC5314 was grown overnight from frozen stocks in yeast extract, peptone, and dextrose (YPD) medium (BD Biosciences) with 100 μg/mL ampicillin (Sigma) in an orbital shaker at 30 °C. Yeast were sub-cultured to ensure early stationary phase. After pelleting and washing with cold PBS, yeast were counted using a LUNA automated cell counter and cell density adjusted in PBS to 100,000 CFUs per 200μl. Mice were injected via lateral tail vein.

### *C. albicans* killing assay

Viable Cy3+ GMPs were sorted 8 hours post transfection and cultured in a humidified incubator at 37°C and 5% CO2 in Stem Span SFEMII supplemented with 1% Penicillin/ Streptomycin and L-Glutamine in addition to 10ng/ml mSCF, 5ng/ml mIL-3 and mIL6 (Peprotech). On day 3 post tiRNA transfection 50K cells were added to a 96-well clear-bottom plate with 5x10^4^ GMPs. *C. albicans* was prepared as described previously and added to each well at a multiplicity of infection of five in 100 μl of complete RPMI (RPMI 1640 with 2 mM L-glutamine, 10% heat-inactivated fetal bovine serum, and 1% penicillin-streptomycin; ThermoFisher Scientific, Waltham, MA). The plate was incubated at 37°C and 5% CO2 for two hours to allow mammalian cell/fungal interaction. Following co-incubation, mammalian cells were lysed with 1% 4x nonidet P40 solution (10 mM Tris HCl, 150 mM sodium chloride, and 5 mM magnesium chloride, titrated to pH 7.5) and wells were supplemented with optimized yeast growth media (MOPS-RPMI; RPMI 1640 containing 2% glucose and 0.165 M MOPS, titrated to pH 7) to support *C. albicans* growth. Then, 10% PrestoBlue Cell Viability Reagent (ThermoFisher Scientific) was added to each well, and the plate was incubated at 37°C with fluorescence measured every 30 minutes for 18 hours by a SpectraMax i3x plate reader (Molecular Devices, Sunnyvale, CA). Fluorescence was plotted versus time, and the time to midcurve (inflection point) was determined using GraphPad Prism 7 software (La Jolla, CA). Healthy human peripheral blood neutrophils were used as a positive control. Cells were isolated using the EasySep Direct Human Neutrophil Isolation Kit (STEMCELL Technologies).

### Statistical analysis

GraphPad PRISM 7 was used to plot data and run statistical analysis. Unpaired student’s t test was used to calculate significance unless indicated otherwise. Sample sizes were based on prior similar work without the use of additional statistical estimations. All measurements were performed on independent biological replicates unless indicated otherwise.

---

Table 1: Abundance (count of reads per million, CPM) of tRNAs in all samples.

Table 2: Fractions of small RNA sequencing reads mapped to genomic elements in all samples.

Table 3: Expression levels of the genes differentially expressed between GMPGFP+ and GMPGFP-cells (> 2-fold change, FDR <0.001)

**Fig. S1:**
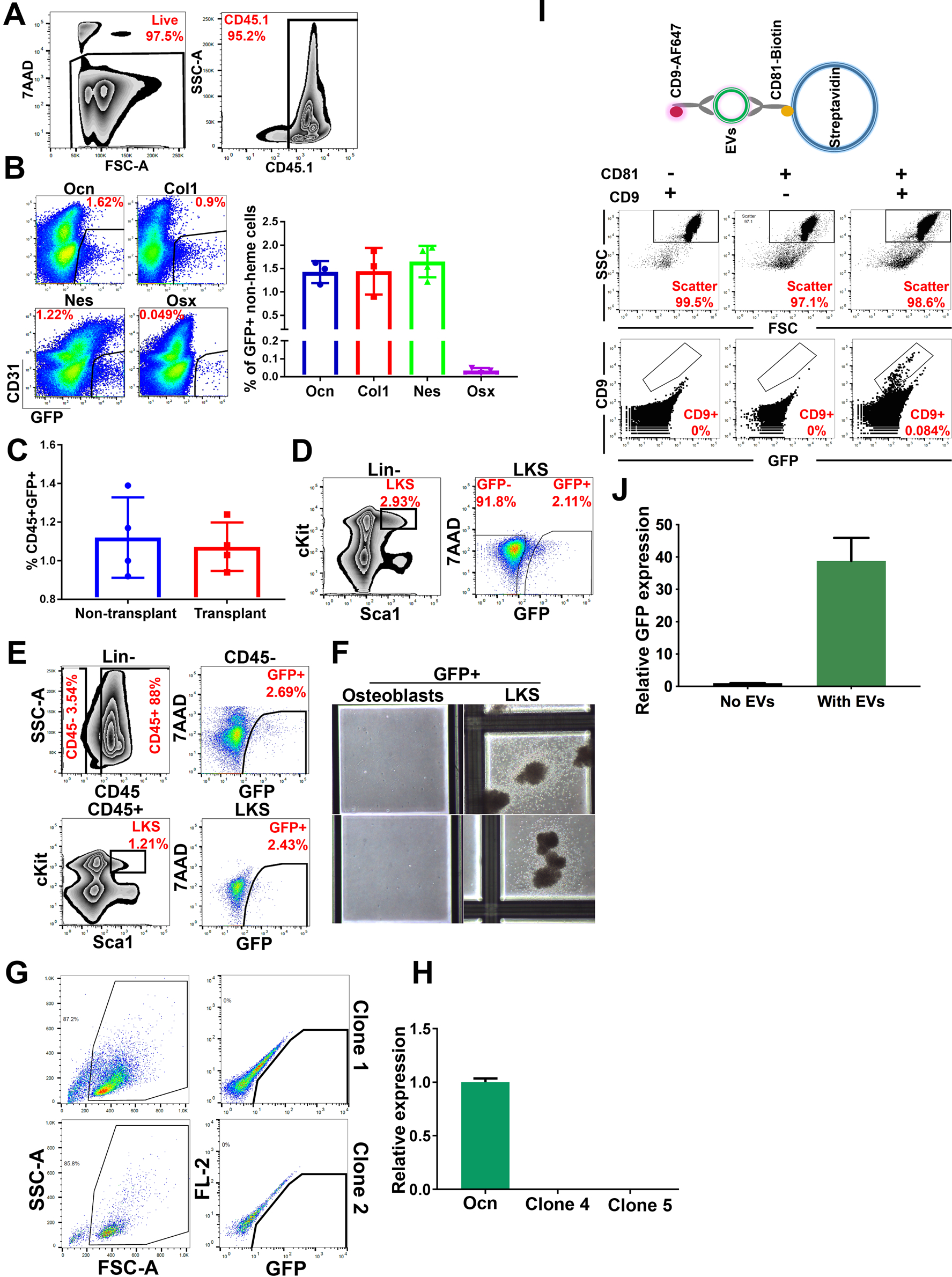
Osteoblast EV transfer is comparable between transplanted and non- transplanted animals. **(A)** Gating strategy for the detection of CD45.1+GFP+ BM cells. **(B)** Frequency of GFP+ mesenchymal cells in non-hematopoietic, non-endothelial bone cells. **(C)** Frequency of CD45+GFP+ BM cells in transplanted and non-transplanted Ocn-GFP animals. **(D)** Gating strategy for LKS^GFP+^ sorted for imaging flow cytometry and confocal microscopy. **(E)** Gating strategy for CD45- GFP+ osteoblasts and CD45+ GFP+ LKS sorted for colony forming assay. **(F)** Hematopoietic colonies in methyl cellulose, images are acquired using 4X objective. **G,H** LKS^GFP+^ methyl cellulose colonies are GFP- **(G)** by flow cytometry and **(H)** qPCR as compared to GFP+ osteoblasts. **(I)** Upper panel, Schematic representation of the flowcytometry assay. Streptavidin beads are coated with EVs bound to biotinylated anti-CD81 and then labeled using anti CD9- AF647. Lower panel, flow cytometry analysis of bead-captured EVs. **(J)** Relative expression of GFP by qPCR in RNA extracted from GMPs cultured with or without Ocn-GFP BM EVs. Data represent three technical replicates. Data in **B**, **C** and **J** is presented as mean ± s.d.

**Fig. S2:**
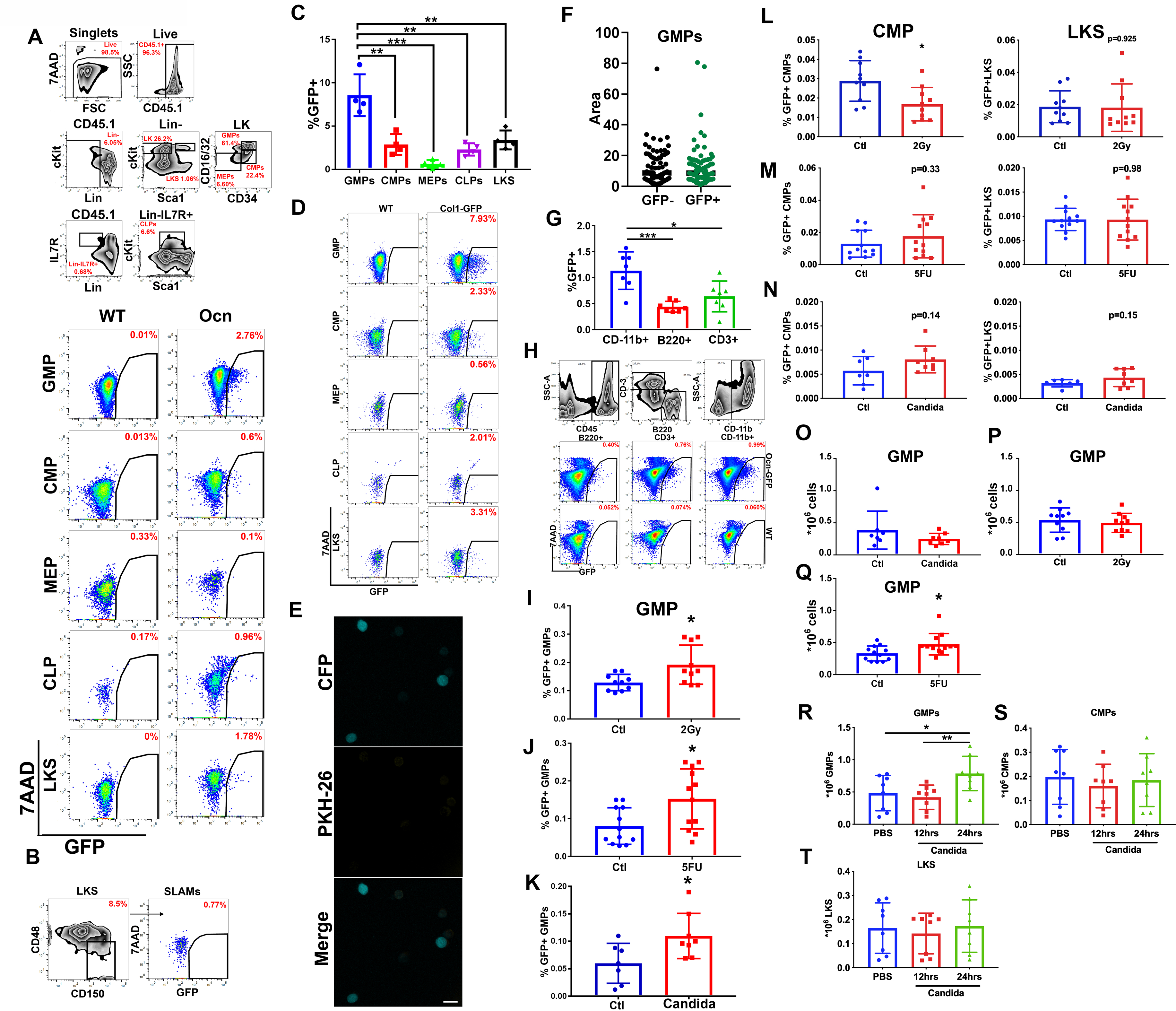
Osteoblastic EVs label immature myeloid progenitors and mature cells in the BM. **(A)** Gating strategy of GFP labeled BM HSPC populations. Parent gates are indicated above the plots (upper) and to the left of the plots (lower). **(B)** Negligible labeling of SLAM HSCs by Ocn-GFP^Topaz^ BM derived EVs. **(C, D)** Labeling of HSPCs by osteoblast derived EVs in the Col1-GFP reporter model. Percentages are of parent gate. Data represent independent biological replicates. **(E)** Maximum projection by confocal imaging of live GMPs demonstrating lack of PKH- 26 labeling in the absence of PKH labeled osteoblasts. Scale bar = 15µm. **(F)** Area of methyl cellulose colonies of GMP^GFP-^ and GMP^GFP+^ as measured by ImageJ. Data represent 6 independent biological replicates with at least 10 colonies representing each replicate. **(G, H)** Osteoblast derived EVs label mature cells in the BM. Percentages are of parent gate. Data represent independent biological replicates. **(I-K)** Frequency of GMP^GFP+^ in total BM mononuclear cells. (**L-N)** Frequency of CMP^GFP+^ and LKS^GFP+^ in total BM live mononuclear cells post-stress **(L)** Irradiation **(M)** 5FU **(N)** *C. albicans* systemic infection. **(O-Q)** Absolute number of GMPs in live mononuclear cells 12 hrs post stress **(R-T)** Absolute number of GMPs, CMPs and LKS in live mononuclear cells 12 and 24 hrs post *C. albicans* infection. Data represent two independent experiments. Data in **C, F, G I-T** is presented as mean ± s.d. *p<0.05, **p<0.01, ***p<0.001.

**Fig. S3:**
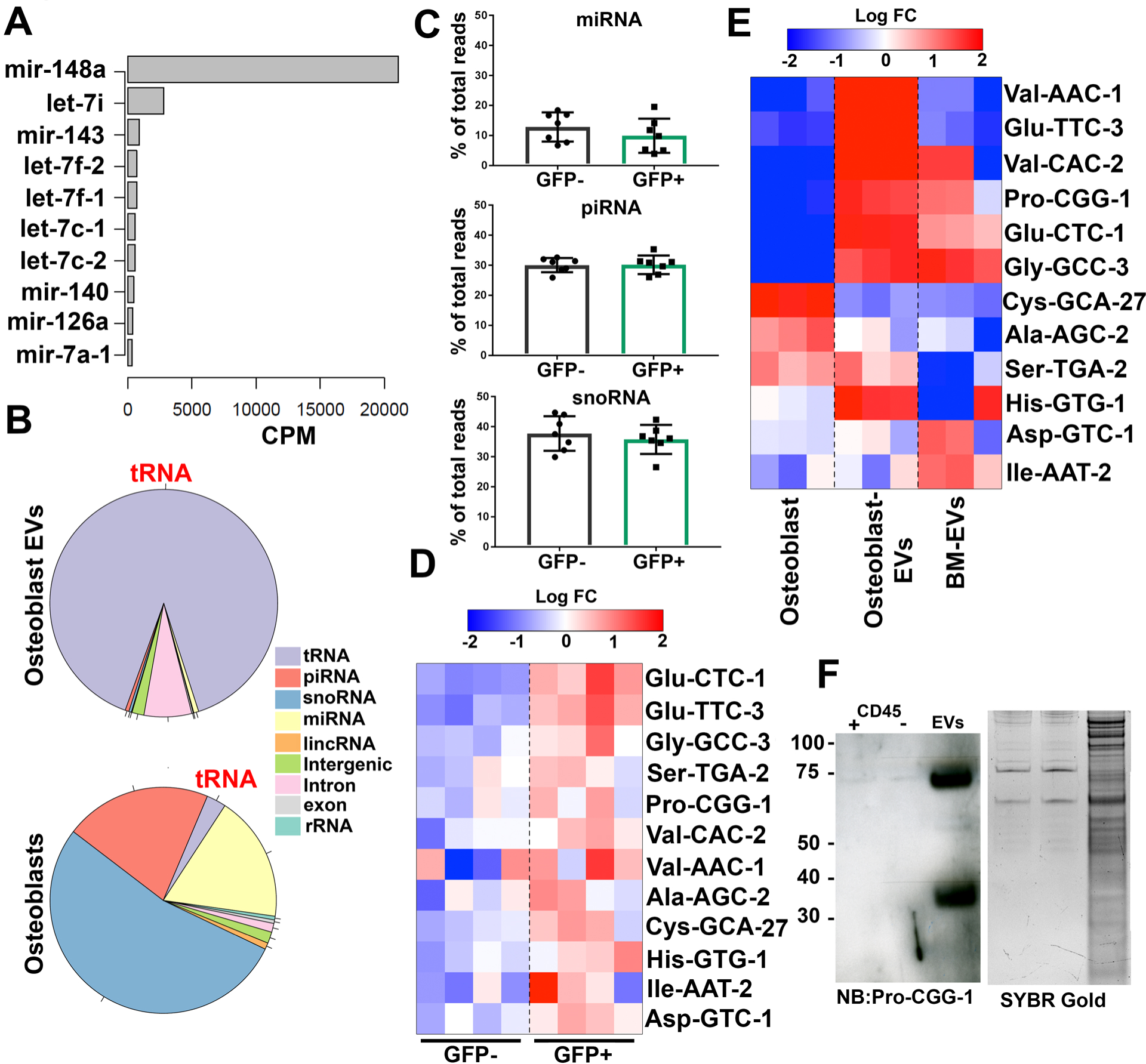
sncRNA in BM and Osteoblast EVs. **(A)** The levels of ten most abundant miRNAs detected in BM EVs, represented as read counts per million (CPM). Data represents three independent biological replicates. **(B)** Fractions of small RNA sequencing reads mapped to genomic elements in osteoblast EVs (upper) and osteoblasts (Lower). Data represents three biological replicates. **(C)** Small RNA fractions in GMP^GFP+^ and GMP^GFP-^. **(D)** Heatmap of tRNAs that are more abundant in GMP^GFP+^ cells > 1.5 fold difference compared to GMP^GFP-^ cells. The levels are shown as relative to the average abundance of a given tRNA across all samples. Data represents one of two independent experiments. **(E)** Heatmap of the tRNA set shown in **D,** comparing the levels of these tRNAs in osteoblasts versus osteoblast EVs and BM EVs. The levels are shown as relative to the average abundance of a given tRNA across all samples. **(F)** Left: northern blot analysis of small RNAs collected from BM CD45+/- cells and BM EVs, Right: SYBR gold stained RNA gel. 500ng Total RNA was loaded.

**Fig. S4:**
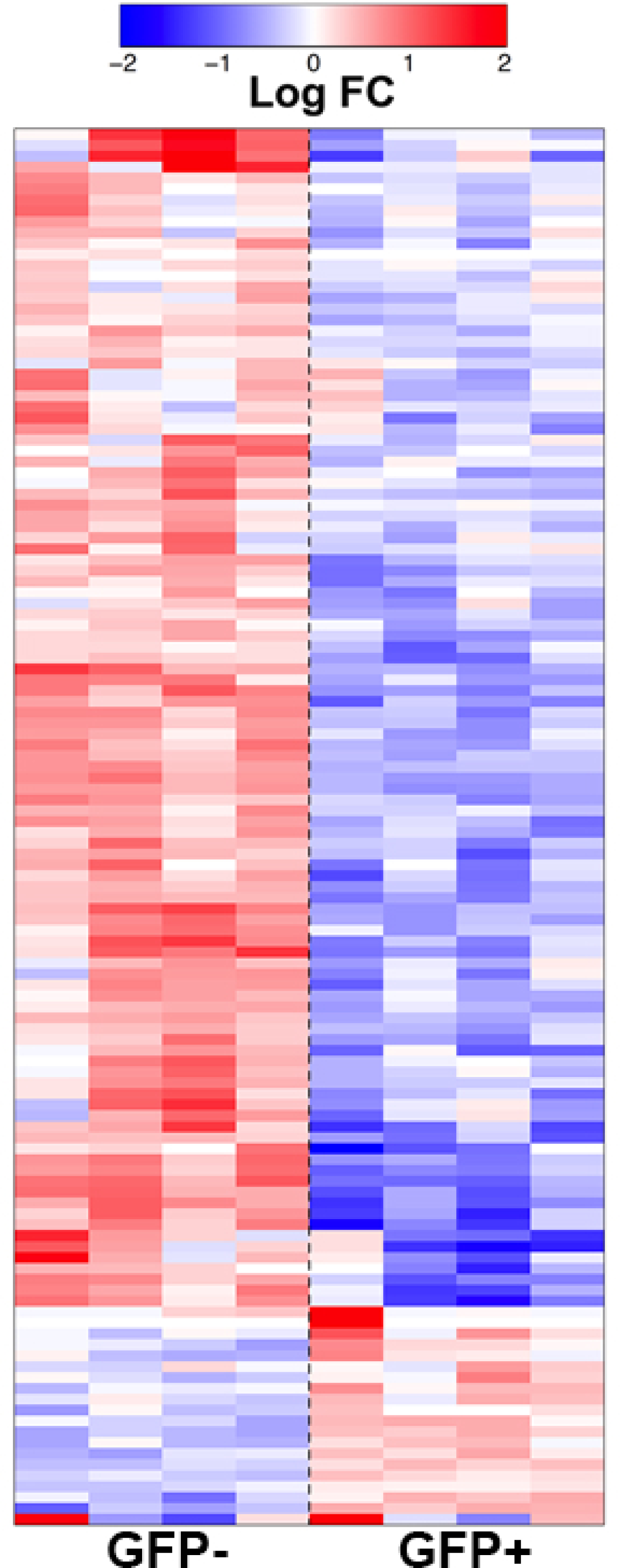
Differential mRNA expression in GMP^GFP+^ vs GMP^GFP-^. Heatmap of expression levels of the genes differentially expressed between GMP^GFP+^ and GMP^GFP-^ cells (> 2-fold change, FDR <0.001). Expression levels are shown as relative to the average for a given gene across all samples.

**Fig. S5:**
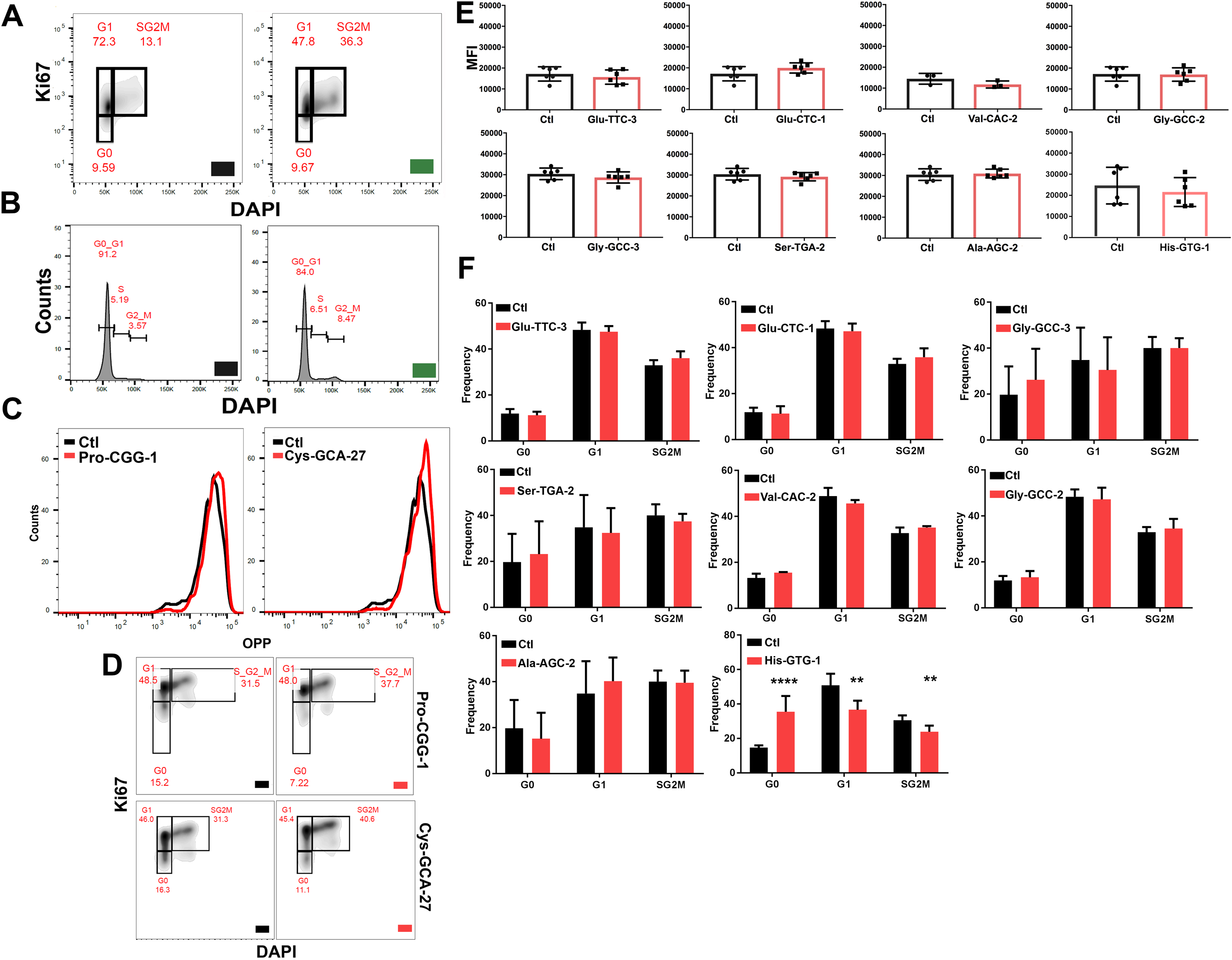
Osteoblastic EVs induce cell proliferation in clonally derived cell lines. **(A)** Gating strategy for cell cycle analysis of GMP^GFP+^ and GMP^GFP-^. **(B)** cell cycle analysis of clonally derived myeloid cell line labeled with EVs. **(C)** OPP uptake and **(D)** gating strategy for cell cycle analysis in Cy3 labeled transfected tiRNA in primary GMPs. **(E)** OPP uptake **(F)** cell cycle analysis of tiRNA transfected GMPs, n=6. Data represent two independent experiments. Data in E and F is presented as mean ± s.d. ** p<0.01, **** p<0.0001.

**Fig. S6:**
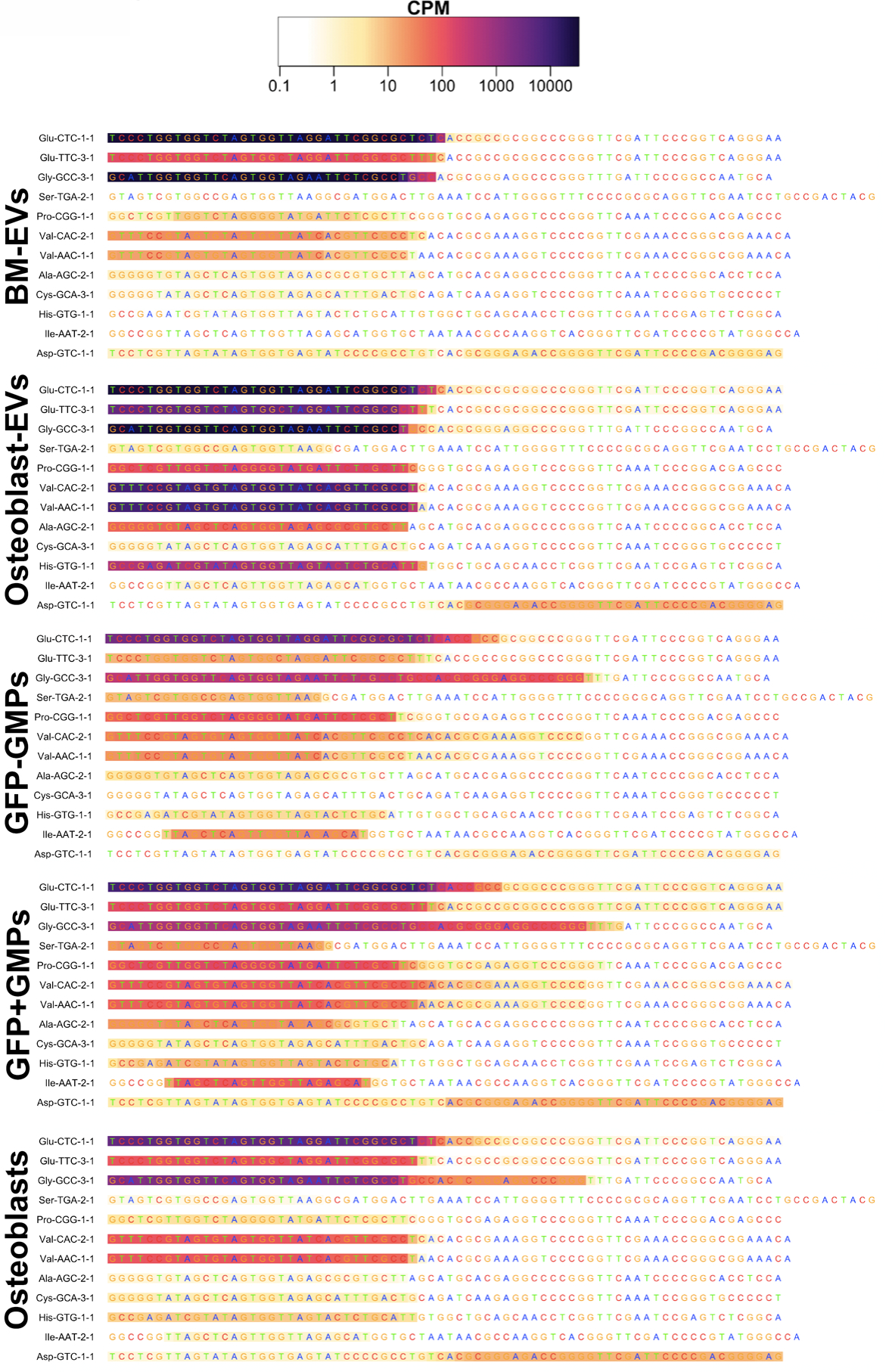
The density of sequencing reads across the length of tRNA sequences. Distributions of density of mapped sequencing reads across the length of tRNA sequences with differential abundance between GMP^GFP+^ vs GMP^GFP-^, shown for BM-EVs, GMP^GFP+^, GMP^GFP-^, osteoblast EVs, and osteoblasts. The sequence of a single tRNA representative is shown for each group of highly similar tRNA species (full groups are listed in table S1). The density of reads (CPM) at each tRNA position is shown by color.

**Fig. S7:**
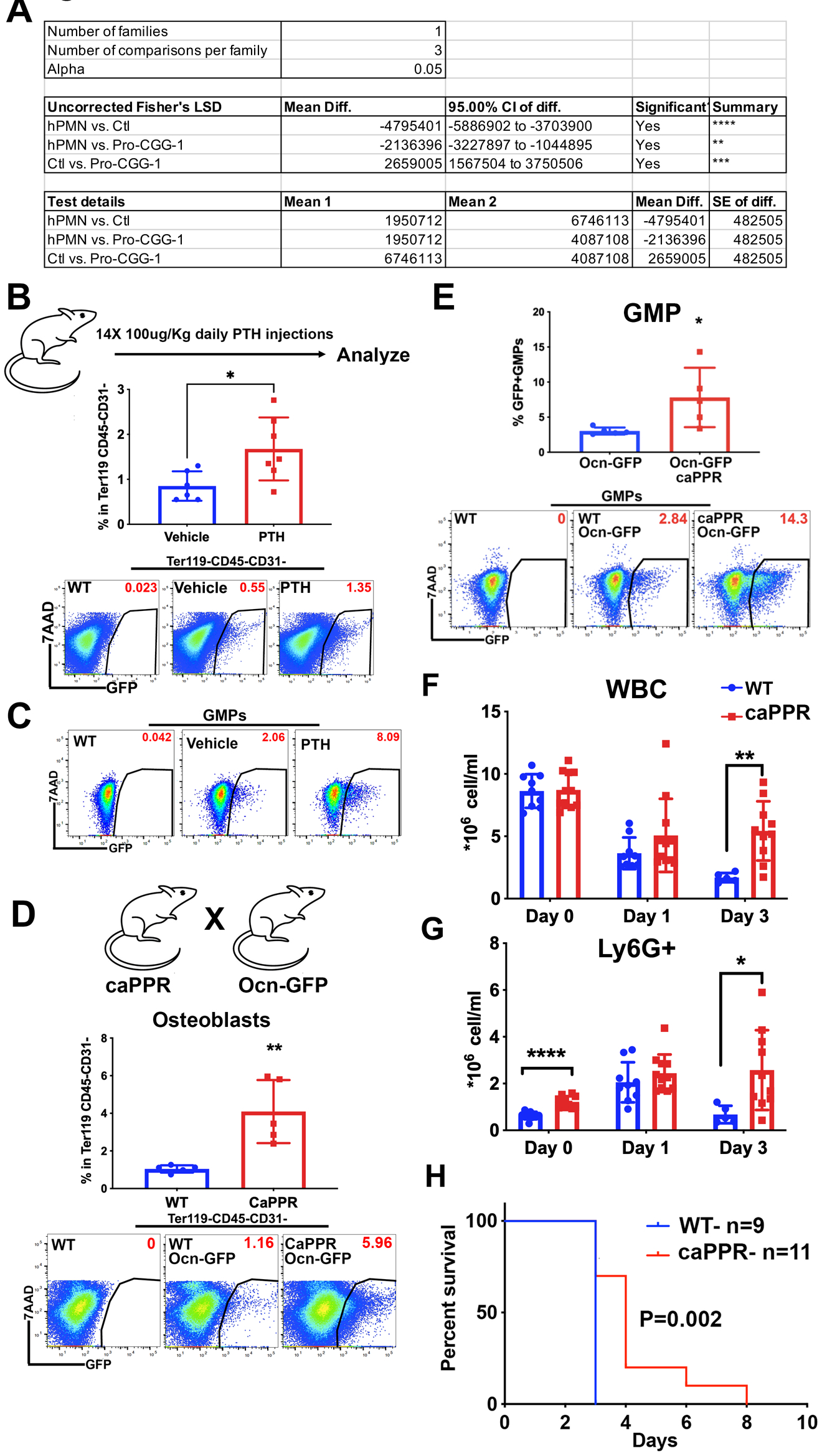
Enhanced transfer of EVs with PTH driven increased osteogenesis. **(A)** One-way ANOVA analysis results of Fig. 5G. **(B)** Frequency of GFP+ osteoblasts (parent gate) within Ter119 CD45- CD31- bone cells 14 days post iPTH treatment. Data represent independent biological replicates from two independent experiments. **(C)** Flow plot demonstrating increased GMP^GFP+^ 14 days post iPTH treatment, percentages are of parent gate. **(D)** Frequency of GFP+ osteoblasts (parent gate) within Ter119 CD45- CD31- bone cells 14 days post iPTH treatment. **(E)** Increased GMP^GFP+^ in caPPR mice. Percentages are of parent gate. **(F)** Peripheral blood WBC **(G)** and neutrophil counts in *C. albicans* infected caPPR mice **(H)** Survival analysis in caPPR mice post *C. albicans* infection. Data represent one of two independent experiments. Data in **D, E, F, G** represent one experiment and is presented as mean ± s.d. *p<0.05, **p<0.01, ****p<0-0001.

